# Bilateral regulation of EGFR activity and local PI dynamics observed with superresolution microscopy

**DOI:** 10.1101/2024.02.01.578337

**Authors:** Mitsuhiro Abe, Masataka Yanagawa, Michio Hiroshima, Toshihide Kobayashi, Yasushi Sako

## Abstract

Anionic lipid molecules, including phosphatidylinositol-4,5-bisphosphate (PI(4,5)P_2_), are implicated in the regulation of epidermal growth factor receptor (EGFR). However, the role of the spatiotemporal dynamics of PI(4,5)P_2_ in the regulation of EGFR activity in living cells is not fully understood, as it is difficult to visualize the local lipid domains around EGFR. In this study, both EGFR and PI(4,5)P_2_ nanodomains in the plasma membrane were visualized using super-resolution single-molecule microscopy. The EGFR and PI(4,5)P_2_ nanodomains aggregated before stimulation with epidermal growth factor (EGF) through transient visits of EGFR to the PI(4,5)P_2_ nanodomains. The degree of coaggregation decreased after EGF stimulation and depended on phospholipase Cγ, the EGFR effector hydrolyzing PI(4,5)P_2_. Artificial reduction in the PI(4,5)P_2_ content of the plasma membrane reduced both the dimerization and autophosphorylation of EGFR after stimulation with EGF. Inhibition of PI(4,5)P_2_ hydrolysis after EGF stimulation decreased phosphorylation of EGFR-Thr654. Thus, EGFR kinase activity and the density of PI(4,5)P_2_ around EGFR molecules were found to be mutually regulated.

## Introduction

Epidermal growth factor receptor (EGFR) is a receptor tyrosine kinase responsible for cell proliferation and differentiation (Nyati et al., 2006; Olayioye et al., 2000). EGFR is subdivided into five regions: a large extracellular region, a single-spanning transmembrane I region, an intracellular juxtamembrane (JM) region, a tyrosine kinase region, and a C-terminal tail region (Lemmon et al., 2014). After EGFR binds to epidermal growth factor (EGF), the extracellular region adopts conformations favoring the dimerization of the transmembrane helices near their N terminus and the dimerization of the JM region (Arkhipov et al., 2013; Ogiso et al., 2002), inducing the formation of asymmetric (active) dimers of the intracellular kinase region (Red Brewer et al., 2009; Thiel and Carpenter, 2007; Zhang et al., 2006). This dimerization results in the phosphorylation of several tyrosine residues in the tail region and the recruitment of intracellular signal proteins, such as growth factor receptor-bound protein 2 (GRB2) and phospholipase Cγ (PLCγ) containing Src homology 2 (SH2) and/or phosphotyrosine-binding regions (Wagner et al., 2013). The JM region of EGFR plays a crucial role in the conformation-dependent coupling of EGFR’s binding to EGF and its activation and dimerization (Endres et al., 2013; Jura et al., 2009b). The JM region comprises a JM-A (N-terminal half) region, which can form an antiparallel helix dimer, and a JM-B (C-terminal half) region, which makes intramolecular contact with the kinase region (Jura et al., 2009a). Both these JM regions contribute to the stable formation of an asymmetric kinase dimer, which is important for kinase activation.

Anionic lipid molecules in the inner leaflet of the plasma membrane are implicated in the dimerization of the JM regions (Hedger et al., 2015; Matsushita et al., 2013; McLaughlin et al., 2005). Multiscale molecular dynamic simulations have suggested that phosphatidylinositol-4,5-bisphosphate (PI(4,5)P_2_) interacts specifically with the basic residues in the JM-A region, stabilizing the JM-A helices in an orientation away from the membrane surface by binding to PI(4,5)P_2_ (Abd Halim et al., 2015; Matsushita et al., 2013). In addition to these *in silico* analyses, our recent *in vitro* study using nanodisc techniques suggested that phosphatidylserine (PS) and PI(4,5)P_2_ stabilize the conformation of the JM-A dimer (Maeda et al., 2018; Maeda et al., 2022). Apart from these *in silico* and *in vitro* studies, the localization and roles of PS and PI(4,5)P_2_ during EGFR activation have not been fully investigated experimentally in living cells due to the difficulty of visualizing the local lipid domains around EGFR in the plasma membrane.

The main difficulty is that the proposed size of the lipid domain is below the diffraction limit of light. Therefore, conventional fluorescence microscopy cannot image the structure in detail. Superresolution imaging techniques that break the diffraction limit have become a powerful tool for visualizing cellular structures with unprecedented resolution (Pujals et al., 2019). Among these techniques, single-molecule localization microscopy (SMLM), which includes photoactivated localization microscopy, fluorescence photoactivation localization microscopy, stochastic optical reconstruction microscopy (STORM), ground-state depletion microscopy followed by individual molecular return, and direct STORM, allows the construction of superresolved images with high-precision localization of individual fluorophores (Rust et al., 2006).

In this study, we visualized both the EGFR and PI(4,5)P_2_ nanodomains in the plasma membrane with SMLM. We found that the PI(4,5)P_2_ nanodomains and EGFR nanoclusters aggregate together before EGF stimulation. After stimulation, the degree of coaggregation decreases, which depends on PLCγ but not on phosphoinositide 3-kinase (PI3K). The local PI(4,5)P_2_ around EGFR stabilizes EGFR dimers to increase EGFR autophosphorylation upon stimulation with EGF. The subsequent hydrolysis of PI(4,5)P_2_ by PLCγ plays a crucial role in the deactivation of EGFR. Our results suggest that autogenous remodeling of the lipid environments regulates EGFR activity.

## Results

### EGF-induced reduction of PI(4,5)P_2_ nanodomains

To visualize the nanodomains of EGFR and the lipids in the plasma membrane simultaneously, we constructed a three-color SMLM analysis workflow (see Methods, Fig. S1). EGFR was fused with rsKame, a slow-switching Dronpa variant (Rosenbloom et al., 2014), at the C-terminus. To exclude the effect of endogenous EGFR, we knocked out endogenous EGFR in HeLa cells, transfected them with EGFR–rsKame, and selected a cell line stably expressing the EGFR–rsKame fusion protein. When we stimulated the selected cell line with EGF, the EGFR–rsKame and extracellular signal-regulated kinase (ERK) in the cell line were phosphorylated to the same levels as in the parental HeLa cells (Fig. S2A and S2B), indicating that the EGFR–rsKame fusion protein functioned normally. To observe PI(4,5)P_2_ with SMLM, the PI(4,5)P_2_-binding peptide PLCδ–PH was fused with a photoactivatable protein, PAmCherry1 (Abe et al., 2012), and was transiently expressed in the cell line. To visualize PS with SMLM, the PS-specific peptide evectin-2 (evt–2)–PH (Uchida et al., 2011), tagged with HaloTag, was transiently expressed in the cell line, and was labeled with a spontaneous blinking dye, HMSiR (Takakura et al., 2017).

After the cells were fixed, sequential three-color SMLM was performed by imaging EGFR–rsKame illuminated with a 488-nm laser; PAmCherry1–PLCδ–PH (hereafter “PAmCherry–PI(4,5)P_2_”) activated with a 405 nm laser and illuminated with a 532 nm laser; and HMSiR–evt–2–PH (hereafter “HMSiR–PS”) illuminated with a 637 nm laser (Fig. S1A). A single-molecule tracking analysis, drift correction, alignment of the color channels, image reconstitution, a multidistance spatial cluster analysis, and a G-function spatial map (G-SMAP) analysis were then performed automatically according to the workflow shown in the Methods section.

EGFR–rsKame, PAmCherry–PI(4,5)P_2_, and HMSiR–PS were distributed nonrandomly in the plasma membrane (Fig. 1A). The G-SMAP analysis revealed that EGFR, PI(4,5)P_2_, and PS formed nanodomains with diameters ranging from <100 nm to several hundred nanometers. We measured the densities and areas of the nanodomains (Fig. 1B). Before EGF stimulation, their size was widely distributed, with a median of 0.0052 ± 0.0002 µm^2^ (∼0.04 µm radius, n = 8 cells) for the EGFR–rsKame nanodomains. Neither the density nor the area of the EGFR–rsKame nanodomains was affected by EGF stimulation (median of 0.0053 ± 0.0001 µm^2^ in area, ∼0.04 radius, n = 10 cells). However, the density of the PAmCherry–PI(4,5)P_2_ nanodomains decreased after EGF stimulation; the median area of PAmCherry–PI(4,5)P_2_ increased from 0.0049 ± 0.0001 µm^2^ (∼0.04 µm radius, n = 8 cells) to 0.0081 ± 0.0004 µm^2^ (∼0.05 µm radius, n = 10 cells, p < 0.001) after EGF stimulation. In particular, the size of smaller PAmCherry–PI(4,5)P_2_ clusters was reduced after EGF stimulation.

**Figure 1.**
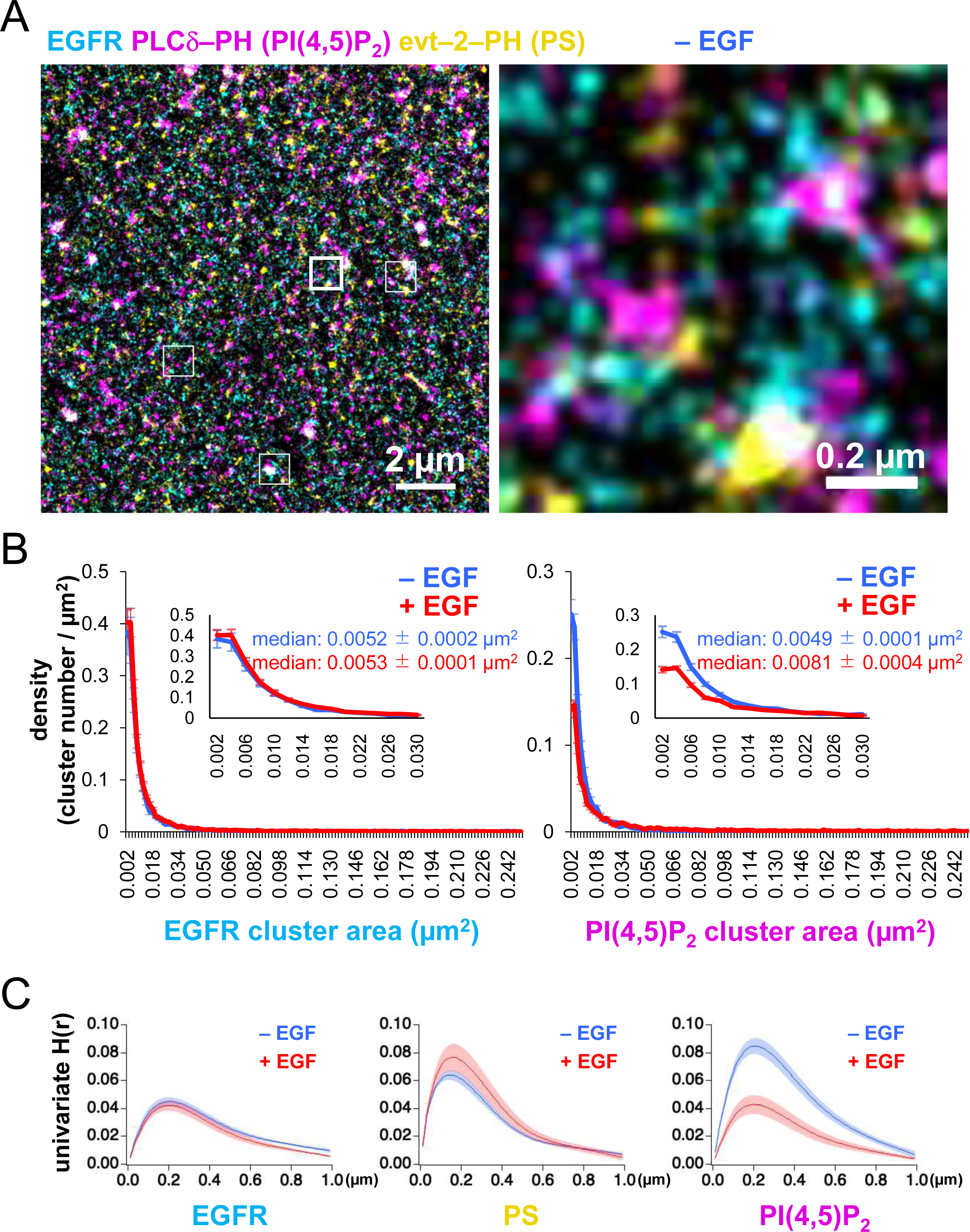
Size and distribution of nanodomains of EGFR, PI(4,5)P_2_, and PS. (**A**) Images of EGFR–rsKame (cyan), PAmCherry–PI(4,5)P_2_ (magenta), and HMSiR–PS (yellow) before EGF stimulation. PAmCherry–PI(4,5)P_2_ and Halo–evt2–PH (Halo–PS) were transiently expressed in a cell line stably expressing EGFR–rsKame. Cells were incubated in serum-free medium in the presence of the HMSiR–Halo ligand overnight and treated with paraformaldehyde and glutaraldehyde. Left, a typical image of 15 × 15 µm. Right, enlarged image of the square region surrounded by a bold line in the left image. Enlarged images of other square regions surrounded by thin lines in the left image are shown in Fig. S2C. (**B**) Distribution of the densities of EGFR (left) and PI(4,5)P_2_ cluster areas (right). Cells were incubated in serum-free medium overnight, stimulated with or without 20 nM EGF for 1 min, and treated with paraformaldehyde and glutaraldehyde. From the SMLM images, the cluster areas and the number of clusters were measured. After the cluster number was normalized to the cell area (density), the cluster density was plotted as a function of the cluster area. Inset, enlarged graphs for the cluster areas of <0.03 µm^2^. Blue and red indicate before and after EGF stimulation, respectively. Data are means ± SEM of eight cells. (**C**) Univariate H(R) values of EGFR–rsKame (left), PAmCherry–PI(4,5)P_2_ (middle), and HMSiR–PS (right) in cells incubated in the absence (blue) or presence (red) of 20 nM EGF for 1 min. Ripley’s univariate H-function was calculated from the SMLM images. Data are means ± SEM of nine (EGFR–rsKame and PAmCherry–PI(4,5)P_2_) or 10 (HMSiR–PS) cells.

We also analyzed the lateral univariate aggregation of the EGFR–rsKame, PAmCherry–PI(4,5)P_2_, and HMSiR–PS molecules, calculating Ripley’s univariate H-function, the derivative of Ripley’s univariate K-function (Fig. 1C) that is frequently used in cluster analyses. The extent of aggregation, the H(r) value, was plotted as a function of the distance between molecules, r, in micrometers. The distance, r, that maximizes the H(r) value (referred to as the R value here) reflects the radius of the area in which the local density of molecules exceeds the random distribution by the largest amount (Kiskowski et al., 2009). The R values of EGFR–rsKame, PAmCherry–PI(4,5)P_2_, and HMSiR–PS were 0.20 ± 0.01 µm (n = 9 cells), 0.21 ± 0.01 µm (n = 9 cells), and 0.14 ± 0.01 µm (n = 10 cells), respectively (Fig. 1C). Therefore, the EGFR, PI(4,5)P_2_, and PS molecules accumulated within membrane domains of similar sizes. The aggregations detected in the H-function analysis (∼0.2 µm; Fig. 1C) were larger than those of the nanodomains estimated with the G-function-based clustering analysis of localization (∼0.04 µm; Fig. 1B), indicating a hierarchical domain structure, that is, the formation of nanodomain clusters in the plasma membrane. The areas of aggregation (domain clusters) were not altered by EGF stimulation for 1 min. The R values for EGFR–rsKame, PAmCherry– PI(4,5)P_2_, and HMSiR–PS were 0.20 ± 0.01 µm (n = 9 cells), 0.20 ± 0.01 µm (n = 9 cells), and 0.15 ± 0.01 µm (n = 9 cells), respectively. The peak H(r) value (referred to as the H(R) value here) indicates the degree of aggregation (Kiskowski et al., 2009). The H(R) value of EGFR–rsKame and HMSiR–PS was not significantly different before and after the addition of EGF (Fig. 1C, left and middle). However, after EGF stimulation, the H(R) value of PAmCherry–PI(4,5)P_2_ decreased from 0.085 ± 0.006 to 0.043 ± 0.007 (n = 9 cells, p < 0.001; Fig. 1C, right).

### Lateral coaggregation of EGFR and PI(4,5)P_2_ decreases after EGF stimulation

EGFR and the lipid nanodomains partly colocalized in the plasma membrane (Figs. 1A and S2C). To estimate the degree of lateral coaggregation of the two different species of molecules, we calculated Ripley’s bivariate H-function (Liu et al., 2023; Zhou et al., 2017). H(r) was plotted as a function of the distance r (Fig. 2A), between the two different species of molecules. The peak value of bivariate H(r) (referred to as the bivariate H(R) value here) for EGFR–rsKame and HMSiR–PS was not significantly altered by EGF stimulation (Fig. 2A, left). The bivariate H(R) value of PAmCherry–PI(4,5)P_2_ and HMSiR–PS also did not change significantly (Fig. 2A, middle). However, after EGF stimulation, the bivariate H(R) values of EGFR–rsKame and PAmCherry–PI(4,5)P_2_ decreased from 0.0126 ± 0.0004 to 0.0066 ± 0.0005 (n = 8 cells, p < 0.001). Therefore, EGF stimulation reduced the coaggregation rate of this pair (Fig. 2A, right).

**Figure 2.**
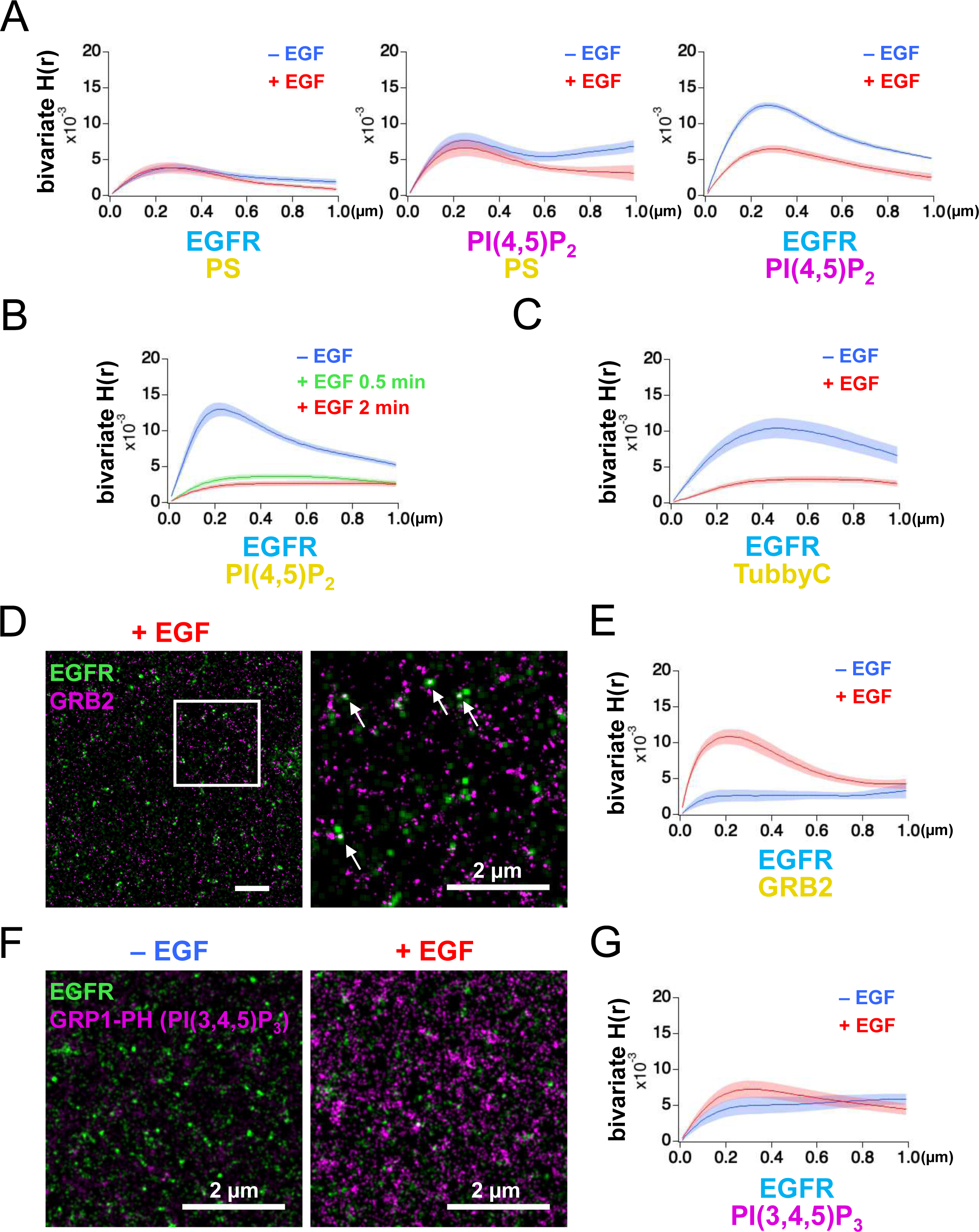
Bivariate H(r) calculated from SLML images. (**A**) Bivariate H(r) value of EGFR–rsKame and HMSiR–PS, PAmCherry–PI(4,5)P_2_ and HMSiR– PS, and EGFR–rsKame and PAmCherry–PI(4,5)P_2_ before (blue) and after incubation with 20 nM EGF for 1 min (red). Cells were prepared as in Fig. 1. Ripley’s bivariate H-function was calculated from the SMLM images. Data are means ± SEM of eight cells. (**B**) Bivariate H(r) value of EGFR–rsKame and HMSiR–PI(4,5)P_2_ before (blue) and after incubation with 20 nM EGF for 0.5 min (green) or 2 min (red). Halo–PI(4,5)P_2_ was transiently expressed in a cell line stably expressing EGFR–rsKame. Cells were incubated in serum-free medium in the presence of the HMSiR–Halo ligand overnight, stimulated with or without 20 nM EGF for the indicated times, and treated with paraformaldehyde and glutaraldehyde. Data are means ± SEM of 17 (without EGF stimulation), 10 (with EGF stimulation for 0.5 min), or 20 cells (with EGF stimulation for 2 min). (**C**) Bivariate H(r) value of EGFR–rsKame and HMSiR–TubbyC before (blue) and after incubation with 20 nM EGF for 2 min (red). Halo–TubbyC was transiently expressed in a cell line stably expressing EGFR–rsKame. Data are means ± SEM of 8 (without EGF stimulation) or 13 cells (with EGF stimulation for 2 min). (**D**) Images of EGFR–rsKame (green) and HMSiR–GRB2 (magenta) after EGF stimulation for 2 min (right). Halo–GRB2 was transiently expressed in a cell line stably expressing EGFR– rsKame. Right, enlarged image of the square region surrounded by a bold line in the left image. Arrows indicate EGFR and GRB2 overlaps in the images. (**E**) Bivariate H(r) value of EGFR–rsKame and HMSiR–GRB2 before (blue) and after incubation with 20 nM EGF for 2 min (red). Halo–GRB2 was transiently expressed in a cell line stably expressing EGFR–rsKame. Data are means ± SEM of 10 cells. (**F**) Images of EGFR–rsKame (green) and PAmCherry–PI(3,4,5)P_3_ (magenta) before (left) and after EGF stimulation for 2 min (right). PAmCherry1–GRP1-PH (PAmCherry–PI(3,4,5)P_3_) was transiently expressed in a cell line stably expressing EGFR–rsKame. (**G**) Bivariate H(r) value of EGFR–rsKame and PAmCherry–PI(3,4,5)P_3_ before (blue) and after incubation with 20 nM EGF for 2 min (red). Data are means ± SEM of 10 (before EGF stimulation) or nine cells (after 20 nM EGF stimulation).

To confirm these results, we performed the following three experiments. First, we replaced PAmCherry–PI(4,5)P_2_ with HMSiR–PI(4,5)P_2_ and observed EGFR–rsKame and HMSiR–PI(4,5)P_2_ with our SMLM system. EGF stimulation for 2 min significantly reduced the bivariate H(R) value of EGFR–rsKame and HMSiR–PI(4,5)P_2_ from 0.0133 ± 0.0010 (n = 17 cells) to 0.0027 ± 0.0003 (n = 20 cells, p < 0.001; Fig. 2B), similar to the results shown in Fig. 2A, right. Even 0.5 min stimulation with EGF induced a decrease in the bivariate H(R) value of EGFR–rsKame and HMSiR–PI(4,5)P_2_. Second, we replaced PLCδ–PH with TubbyC, another biosensor for PI(4,5)P_2_, because the PI(4,5)P_2_ biosensors show differences in mobility in the plasma membrane (Pacheco et al., 2022). Similar to the results in Fig. 2B, EGF stimulation significantly reduced the bivariate H(R) value of EGFR–rsKame and HMSiR–TubbyC from 0.0108 ± 0.0013 (n = 8 cells) to 0.0038 ± 0.0004 (n = 13 cells, p < 0.001; Fig. 2C). Third, we observed EGFR–rsKame and HMSiR– GRB2 with our SMLM system and measured the bivariate H(R) value. We found several EGFR and GRB2 overlaps in the SMLM images after EGF stimulation for 2 min (Fig. 2D).

EGF stimulation increased the bivariate H(R) value of EGFR–rsKame and HMSiR–GRB2 from 0.0027 ± 0.0008 (n = 10 cells) to 0.0110 ± 0.0010 (n = 10 cells, p < 0.001; Fig. 2E). These values are consistent with the finding that cytoplasmic GRB2 is recruited to phosphorylated EGFR after EGF stimulation (Yoshizawa et al., 2021). These three experiments support the idea that the H-function analysis of the SMLM data reflects the degree of coaggregation, and that EGF stimulation reduced the colocalization rate of EGFR and PI(4,5)P_2_.

Because PI(4,5)P_2_ and PI(3,4,5)P_3_ have similar characteristics but play different roles in the plasma membrane (Insall and Weiner, 2001), we also examined the coaggregation of EGFR and PI(3,4,5)P_3_ before and after EGF stimulation. PAmCherry1– GRP1-PH (hereafter “PAmCherry–PI(3,4,5)P_3_”), which specifically interacts with PI(3,4,5)P_3_, was expressed in the EGFR–rsKame stable cell line (Fig. 2F) and the bivariate H(r) values of EGFR–rsKame and PamCherry–PI(3,4,5)P_3_ were measured (Fig. 2G). The density of PAmCherry–PI(3,4,5)P_3_ was small with a low bivariate H(R) value before EGF stimulation. After EGF stimulation, the bivariate H(R) value was still low (Fig. 2G, red), although, consistent with a previous report (Malek et al., 2017), the number of PI(3,4,5)P_3_ nanodomains stained with PAmCherry–PI(3,4,5)P_3_ increased (Fig. 2F). Thus, the nanodomains of PI(4,5)P_2_ and PI(3,4,5)P_3_ displayed distinct characteristics in their interactions with EGFR before and after EGF stimulation.

### SMT analysis of EGFR and PI(4,5)P_2_

Because the cells used in SMLM were artificially fixed with paraformaldehyde and glutaraldehyde, we examined the colocalization of EGFR and PI(4,5)P_2_ in living cells using single-molecule tracking (SMT) analysis (Figs. 3 and S3). For SMT analysis, we tagged the cytoplasmic tail of EGFR with enhanced green fluorescent protein (EGFR–GFP) (Hiroshima et al., 2018; Yasui et al., 2018) and expressed it in EGFR-knockout (KO) HeLa cells at low levels. EGFR–GFP in the cells prepared for SMT analysis was phosphorylated to the same level as in the parental HeLa cells after EGF stimulation (Fig. S3A), suggesting that EGFR–GFP functioned normally. After the EGFR–GFP and Halo–PI(4,5)P_2_ fusion proteins were expressed in EGFR-KO HeLa cells, Halo–PI(4,5)P_2_ was labeled with Janelia Fluor 549 (JF549). These proteins were observed with dual-color total internal reflection fluorescence for SMT analysis (Fig. 3A). The movements of the fluorescent particles were classified into three motional modes (immobile, slow-mobile, and fast-mobile) according to the Variational Bayesian-Hidden Markov Model (Yanagawa and Sako, 2021; Yasui et al., 2018). The proportion of EGFR in the immobile fraction increased and the ratio in the fast-mobile fraction decreased after EGF stimulation (Fig. 3B) at the same level as that reported previously for EGFR–GFP (Hiroshima et al., 2018; Yasui et al., 2018). The proportion of PI(4,5)P_2_ in each fraction did not change significantly after EGF stimulation. The mean square displacement per unit time (MSD-Δt) of the lateral movements indicated that the diffusion coefficient of EGFR decreased after EGF stimulation, whereas that of PI(4,5)P_2_ remained unchanged (Figs. 3C and S3B). All diffusion modes displayed subdiffusion. The confinement length of EGFR was also reduced after EGF stimulation, whereas that of PI(4,5)P_2_ was not (Fig. S3C).

**Figure 3.**
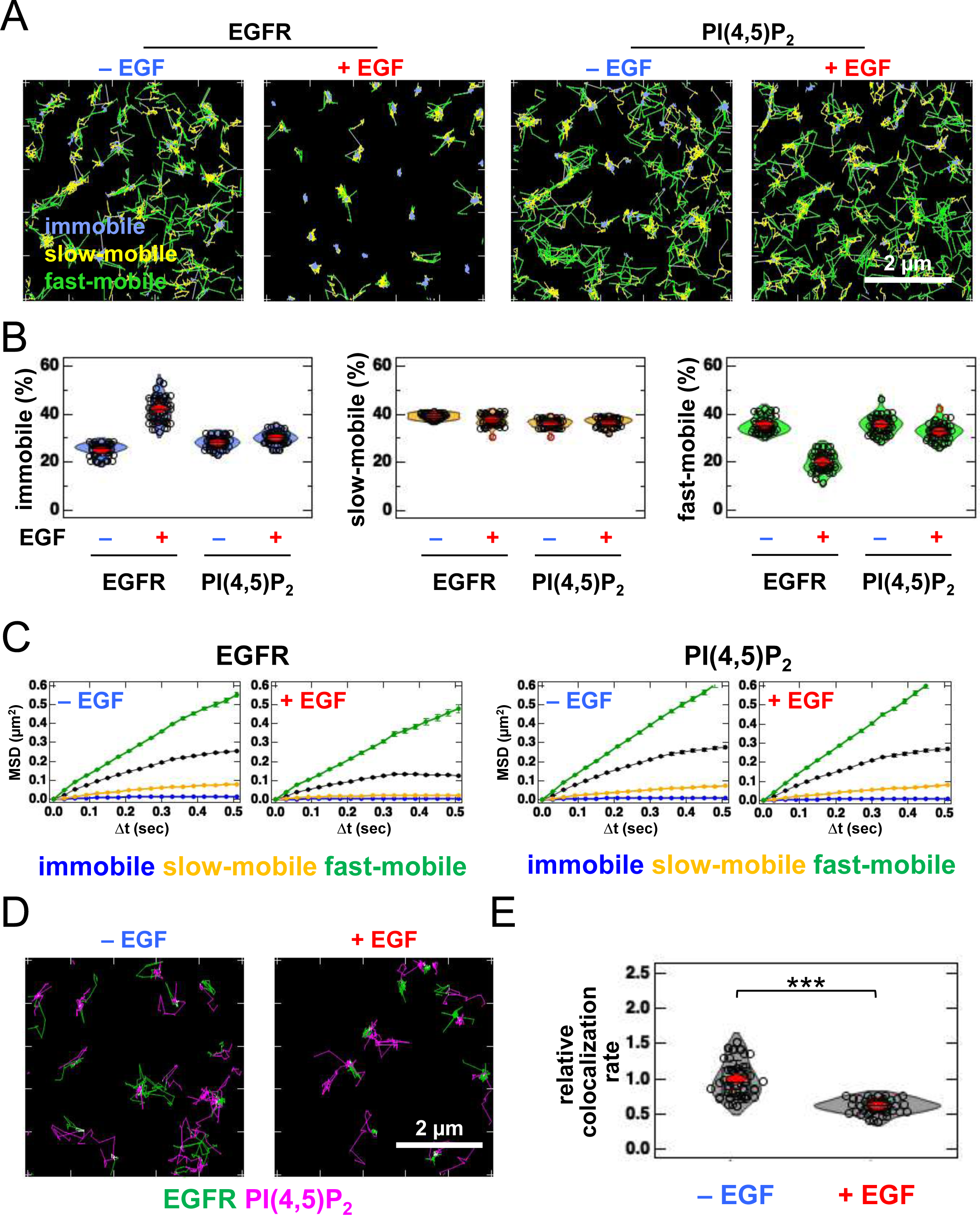
Lateral coaggregation of EGFR and PI(4,5)P_2_ decreases after EGF stimulation. (**A**) Trajectories of EGFR–GFP (left) and JF549–PI(4,5)P_2_ (right) over 6 s. EGFR–GFP and Halo–PLCδ–PH (Halo–PI(4,5)P_2_) were transiently expressed in EGFR-KO HeLa cells. After the cells were incubated in serum-free medium overnight, Halo–PI(4,5)P_2_ was stained with JF549– Halo ligand. EGFR–GFP and JF549–PI(4,5)P_2_ were observed in 30 cells at a time resolution of 30 ms for 6 s before EGF stimulation. After stimulation with 20 nM EGF, EGFR–GFP and JF549–PI(4,5)P_2_ were observed in the same cells between 1 and 5 min after the addition of EGF. SMT analysis was fractionated into the immobile (blue), slow-mobile (yellow), and fast-mobile (green) fractions. (**B**) Proportions of immobile (blue), slow-mobile (yellow), and fast-mobile (green) fractions. (**C**) MSD-Δt plots of the trajectories of EGFR–EGFP and JF549–PI(4,5)P_2_. (**D**) Trajectories of EGFR–GFP (green), JF549–PI(4,5)P_2_ (magenta), and colocalization (white) during 6 s before (left) and after incubation with 20 nM EGF (right) in the same cell. (**E**) Relative colocalization rates of EGFR–GFP and JF549–PI(4,5)P_2_ before and after incubation with 20 nM EGF. To consider the differences in expression levels among cells, the colocalization rate was divided by the densities of EGFR–GFP and JF549–PI(4,5)P_2_ and normalized to the mean value obtained before EGF stimulation. Violin plots show the mean value and distribution of 30 cells. ***p < 0.001 on Welch’s *t* test.

Next, we examined the colocalization of EGFR and PI(4,5)P_2_. We found partial and transient overlaps in the trajectory images of EGFR and PI(4,5)P_2_, before and after EGF stimulation (Fig. 3D; Movie 1). To consider the differences in expression levels among cells, we normalized the original colocalization rate (Yanagawa and Sako, 2021) to the densities of EGFR and PI(4,5)P_2_ (i.e., particle number/cell area). The normalized data show that the colocalization of EGFR and PI(4,5)P_2_ decreased after EGF stimulation (Fig. 3E).

Among the three motional modes, the colocalization rate in the slow-mobile and fast-mobile fractions decreased after EGF stimulation, whereas the rate in the immobile fraction did not change significantly (Fig. S3D). We also examined the colocalization of EGFR and TubbyC (Fig. S3E). The colocalization rate of EGFR and TubbyC were similar to those of EGFR and PLCδ–PH before EGF stimulation and decreased to rate comparable to those of EGFR and PLCδ–PH after EGF stimulation (Fig. S3F). The colocalization rate of EGFR and PI(4,5)P_2_ decreased significantly by 0.5 min after EGF stimulation and remained low for at least 5 min (Fig. S4A). Under the same experimental conditions, the colocalization rate of EGFR and GRB2 increased by 0.5 min after EGF stimulation and remained high for at least 5 min (Fig. S4B). These results suggest that the binding of EGF to EGFR and the subsequent reaction occurred within 0.5 min under our experimental conditions. The transient colocalization of EGFR and PI(4,5)P_2_ observed in SMT analysis was consistent with the small spatial overlap of the nanodomains of these two molecular species in the SMLM images.

### PI(4,5)P_2_ is important for stabilizing the EGFR dimer

*In silico* and *in vitro* analyses have suggested that PI(4,5)P_2_ is involved in the dimerization of EGFR (Abd Halim et al., 2015; Maeda et al., 2018; Maeda et al., 2022; Matsushita et al., 2013). The aggregation of PI(4,5)P_2_ with EGFR in the plasma membrane may help to stabilize the dimers of EGFR after its association with EGF. To test this, we reduced the content of PI(4,5)P_2_ by expressing synaptojanin (SYNJ), a phosphatase that hydrolyzes PI(4,5)P_2_, and measured the size of the EGFR–Halo oligomer in EGFR-null CHO-K1 and EGFR-KO HeLa cells by SMT analysis (Figs. 4A and S5A). The cytoplasmic expression of SYNJ, decreases the amount of PI(4,5)P_2_ in the plasma membrane (Field et al., 2005). EGFR–Halo was labeled with SaraFluor 650T (SF650). In the immobile fraction of control CHO-K1 cells, 55.1 ± 2.6% (n = 40 cells) of the EGFR particles were monomers, whereas 29.7 ± 0.8% (n = 40 cells) were preformed dimers before EGF stimulation (Figs. 4A, S5B, and S5C). After EGF stimulation, the monomers decreased to 38.8 ± 4.1% (n = 40 cells) and the dimers increased to 36.9 ± 1.8% (n = 40 cells) in the control cells. The slow- and fast-mobile fractions showed little change in EGFR oligomer size. However, in the immobile fraction of SYNJ-expressing cells, the monomers decreased slightly even after EGF stimulation (57.6 ± 2.7% before and 50.3 ± 3.4% after EGF stimulation, n = 40 cells). Similar suppression of EGFR dimers by SYNJ expression was observed in the immobile fraction of EGFR-KO HeLa cells (Fig. S5A). The mean density and mean intensity of EGFR–SF650 in the SYNJ-expressing cells before EGF stimulation were similar to those in the control cells (Fig. S5D and S5E), indicating that the expression levels of EGFR–SF650 were not significantly altered in SYNJ-expressing cells. These results suggest that PI(4,5)P_2_ is important for stabilizing EGFR dimers after EGF stimulation.

**Figure 4.**
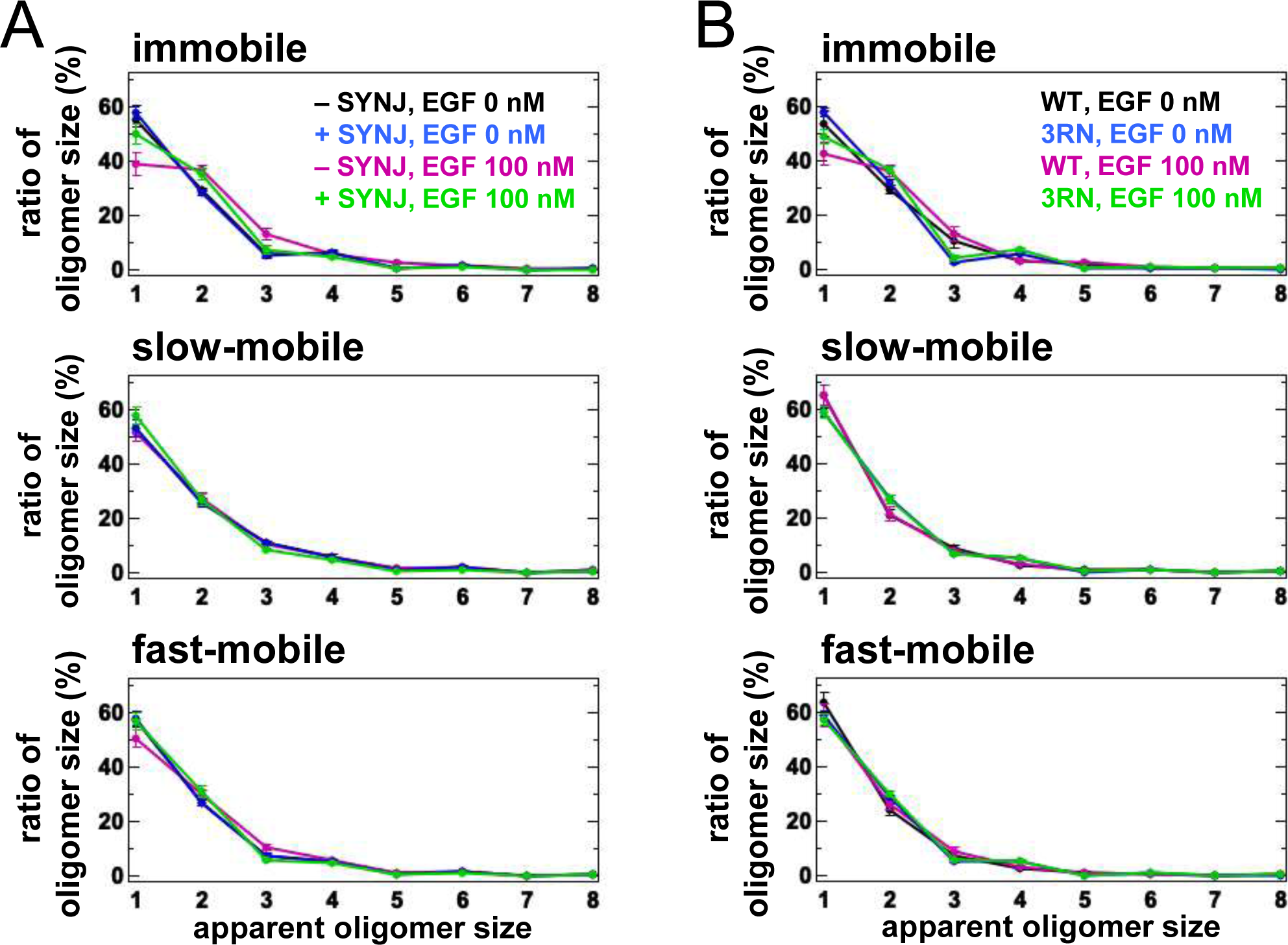
SMT analysis of EGFR under PI(4,5)P_2_-interaction-depleted conditions. (**A**) SMT analysis of EGFR–SF650 in control and SYNJ-expressing cells. Only EGFR–Halo or both EGFR–Halo and GFP–SYNJ were transiently expressed in CHO-K1 cells. After cells were incubated in serum-free medium overnight, EGFR–Halo was stained with the SF650–Halo ligand. EGFR–SF650 was observed in control (n = 40) and SYNJ-expressing cells (n = 40) at a time resolution of 30 ms before EGF stimulation. Following stimulation with 100 nM EGF, EGFR–SF650 was observed in the same cells between 1 and 6 min after the addition of EGF. (**B**) SMT analysis of EGFR(WT)–SF650 and EGFR(3RN)–SF650. EGFR–Halo or EGFR(3RN)–Halo was transiently expressed in CHO-K1 cells. After the cells were incubated in serum-free medium overnight, EGFR–Halo was stained with the SF650–Halo ligand. EGFR(WT)–SF650 (n = 40) and EGFR(3RN)–SF650 (n = 45) were observed as described in (A). Data are means ± SEM.

Molecular dynamics simulations have shown that the conversion of three arginine residues to asparagine residues in the EGFR JM region (EGFR(3RN)) abolishes the interaction between EGFR and PI(4,5)P_2_ (Abd Halim et al., 2015). Therefore, we examined the oligomer size of EGFR(3RN) in CHO-K1 cells before and after EGF stimulation (Figs. 4B, S6A, and S6B). Similar to oligomer size of EGFR(WT), that of EGFR(3RN) showed little change in the slow- and fast-mobile fractions. The monomer in the immobile fraction constituted 48.9 ± 2.5% of EGFR(3RN) (n = 45 cells) after EGF stimulation (Fig. 4B), which was similar to that in the SYNJ-expressing cells (Fig. 4A). The relative intensity of Cy5–EGF bound to EGFR did not differ significantly between EGFR(WT) and EGFR(3RN), suggesting that the 3RN mutation does not affect the binding of EGFR to EGF (Fig. S6C and S6D). These results suggest that the interaction of EGFR with PI(4,5)P_2_ is important for the dimerization of EGFR.

In SMT, molecular-level interactions of dimers cannot be detectable. Therefore, we confirmed these direct interactions with a biochemical analysis using a crosslinker (Fig. 5A and 5B). Cells were treated with 0.2 or 20 nM EGF, which is below or above the dissociation constant of 2–6 nM (Sugiyama et al., 2023), respectively, and cell surface EGFR was crosslinked with EGFR using bis(sulfosuccinimidyl)suberate. The dimer fraction of EGFR was small before EGF stimulation but increased after stimulation with 0.2 or 20 nM EGF. Under both conditions, the expression of SYNJ reduced the amount of EGFR dimer (Fig. 5A and 5B), suggesting that PI(4,5)P_2_ is involved in stabilizing the dimerization of EGFR.

**Figure 5.**
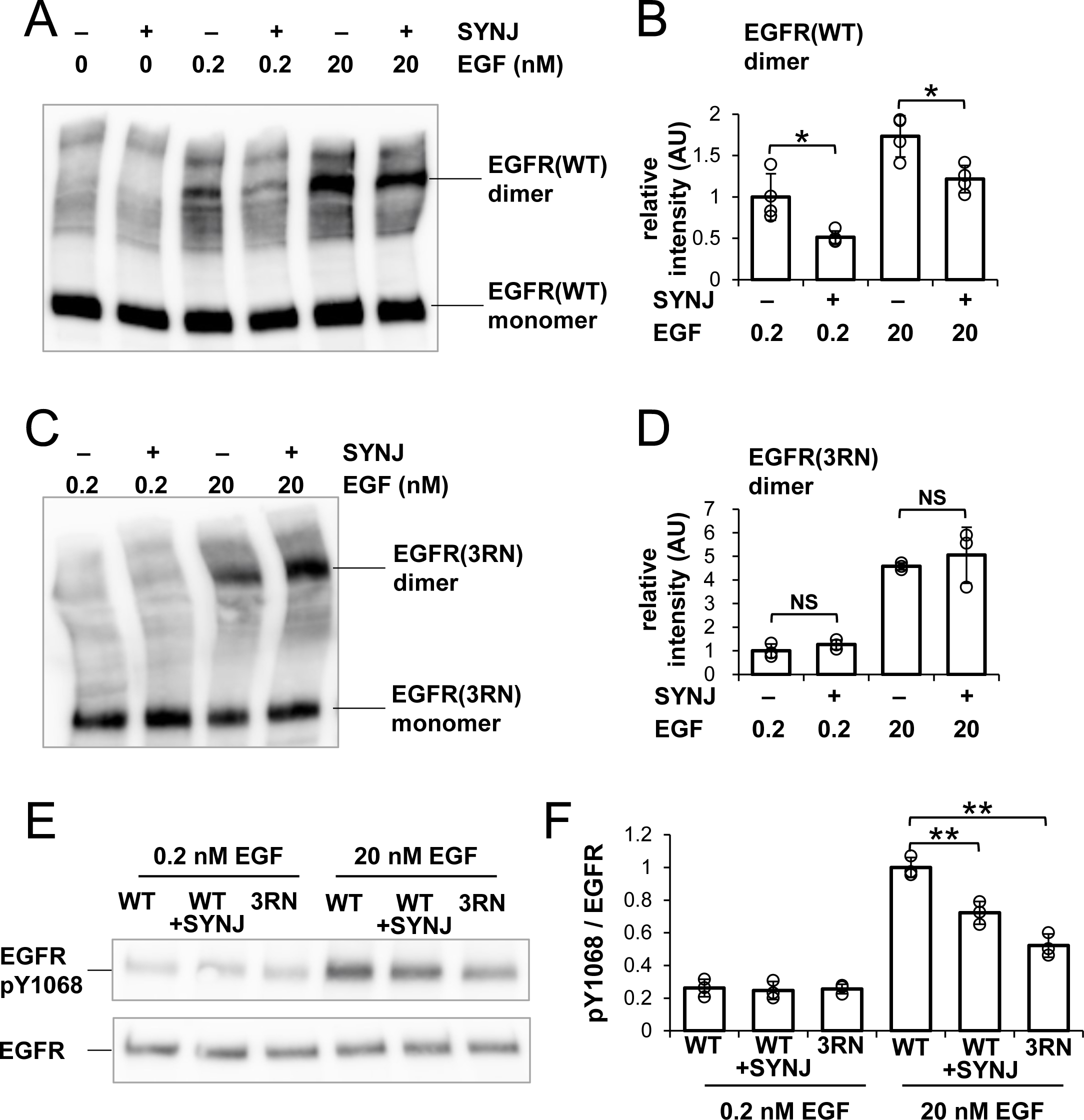
Biochemical analysis of EGFR under PI(4,5)P_2_-depleted conditions. (**A**) Western blotting analysis of crosslinked EGFR. After EGFR-KO cells were transfected with only EGFR–Halo or both EGFR–Halo and GFP–SYNJ, the cells were incubated in serum-free medium overnight. The cells were then stimulated with 0, 0.2, or 20 nM EGF, and treated with a crosslinker for 1 h. EGFR–Halo was detected with an anti-EGFR antibody. (**B**) Amounts of EGFR–Halo dimer. Relative intensity was normalized to the mean intensity of lane #3 in (A). Data are means ± SD of four experiments. (**C**) Western blotting analysis of crosslinked EGFR(3RN). After EGFR-KO cells were transfected with only EGFR(3RN)–Halo or both EGFR(3RN)–Halo and GFP–SYNJ, the cells were incubated in serum-free medium overnight. The cells were then stimulated with 0.2 or 20 nM EGF, and treated with a crosslinker for 1 h. EGFR(3RN)–Halo was detected with an anti-EGFR antibody. (**D**) Amounts of EGFR(3RN)–Halo dimer. Relative intensity was normalized to the mean intensity of lane #1 in (C). Data are means ± SD of three experiments. (**E**) Western blotting analysis of phosphorylated EGFR and total EGFR. After EGFR-KO cells were transfected with EGFR(WT)–Halo, EGFR(WT)–Halo and GFP–SYNJ, or EGFR(3RN)– Halo, the cells were incubated in serum-free medium overnight and stimulated with 0.2 nM or 20 nM EGF for 2 min. Phospho-EGFR and total EGFR were detected with anti-pY1068 EGFR and EGFR, respectively. (**F**) Ratio of phosphorylated-Tyr1068 EGFR/total EGFR. The ratio was normalized to the mean value of lane #4 in (E). Data are means ± SD of three experiments. *p < 0.05, **p < 0.01, NS (not significant) on Welch’s *t* test.

Because the expression of SYNJ does not completely abolish PI(4,5)P_2_ in the cell (Field et al., 2005), we could not determine whether PI(4,5)P_2_ is essential for the dimerization of EGFR. To test this, we examined whether EGFR(3RN) forms dimers after EGF stimulation (Fig. 5C and 5D). After stimulation with 0.2 or 20 nM EGF, the amount of EGFR(3RN) dimers increased. If the interaction with PI(4,5)P_2_ was not completely abolished in EGFR(3RN)-expressing cells, the amount of EGFR(3RN) dimers should decrease when PI(4,5)P_2_ was reduced. There was no additive effect on the dimerization of EGFR(3RN) when SYNJ was expressed (Fig. 5C and 5D), suggesting that the interaction of EGFR(3RN) with PI(4,5)P_2_ was rigorously blocked. Therefore, PI(4,5)P_2_ is not essential for, but positively regulates EGFR dimerization.

The dimerization of EGFR leads to the phosphorylation of tyrosine residues in its tail region (Wagner et al., 2013). We examined whether the reduced dimerization of EGFR under PI(4,5)P_2_-interaction-depleted conditions resulted in defective phosphorylation of EGFR (Fig. 5E and 5F). The measurement of Tyr1068-phosphorylated EGFR after stimulation with 20 nM EGF indicated that the phosphorylation of EGFR decreased when SYNJ was overexpressed in cells. We also found that the Tyr1068 phosphorylation in EGFR(3RN) decreased relative to that in EGFR(WT) after stimulation with 20 nM EGF. However, after simulation with 0.2 nM EGF, no significant difference in the phosphorylation of EGFR was observed between the cells as EGFR was less phosphorylated under the current experimental conditions. These results confirm that PI(4,5)P_2_ is important for not only the dimerization of EGFR but also its subsequent phosphorylation.

### Extent of reduced coaggregation depends on PLC**_γ_** but not on PI3K

A biochemical analysis showed that the amount of PI(4,5)P_2_ is transiently reduced after EGF stimulation in cells (Malek et al., 2017). Two kinds of enzymes are activated by EGFR and reduce the amount of PI(4,5)P_2_ after EGF stimulation (Wells, 1999). First, PI3K phosphorylates PI(4,5)P_2_ to produce PI(3,4,5)P_3_ through the heterodimerization of ERBB3 with EGFR (Hellyer et al., 1998). Second, phosphoinositide-specific PLCγ hydrolyzes PI(4,5)P_2_ to produce diacylglycerol (DAG) and inositol trisphosphate (IP_3_) through its direct physical association with EGFR via the SH2 regions of PLCγ (Carpenter and Ji, 1999). We investigated the protein responsible for the dissociation of EGFR from the PI(4,5)P_2_ nanodomains after EGF stimulation. We inhibited either PI3K or PLCγ and measured the bivariate H(r) value of EGFR and PI(4,5)P_2_ after EGF stimulation.

The addition of wortmannin, a PI3K inhibitor (Yano et al., 1993), diminished the PI3K-dependent phosphorylation of AKT after EGF stimulation (Fig. 6A). The expression of a dominant negative fragment of PLCγ containing the SH2, SH3, and PLC-inhibitory regions (referred to as “DN PLCγ” here) inhibits PLCγ activity (Banan et al., 2001; Homma and Takenawa, 1992). We found that the expression of DN PLCγ reduced DAG production after EGF stimulation compared with that in the non-DN-PLCγ-expressing control cells, indicating that the expressed fragment inhibited PLCγ activity (Fig. 6B).

**Figure 6.**
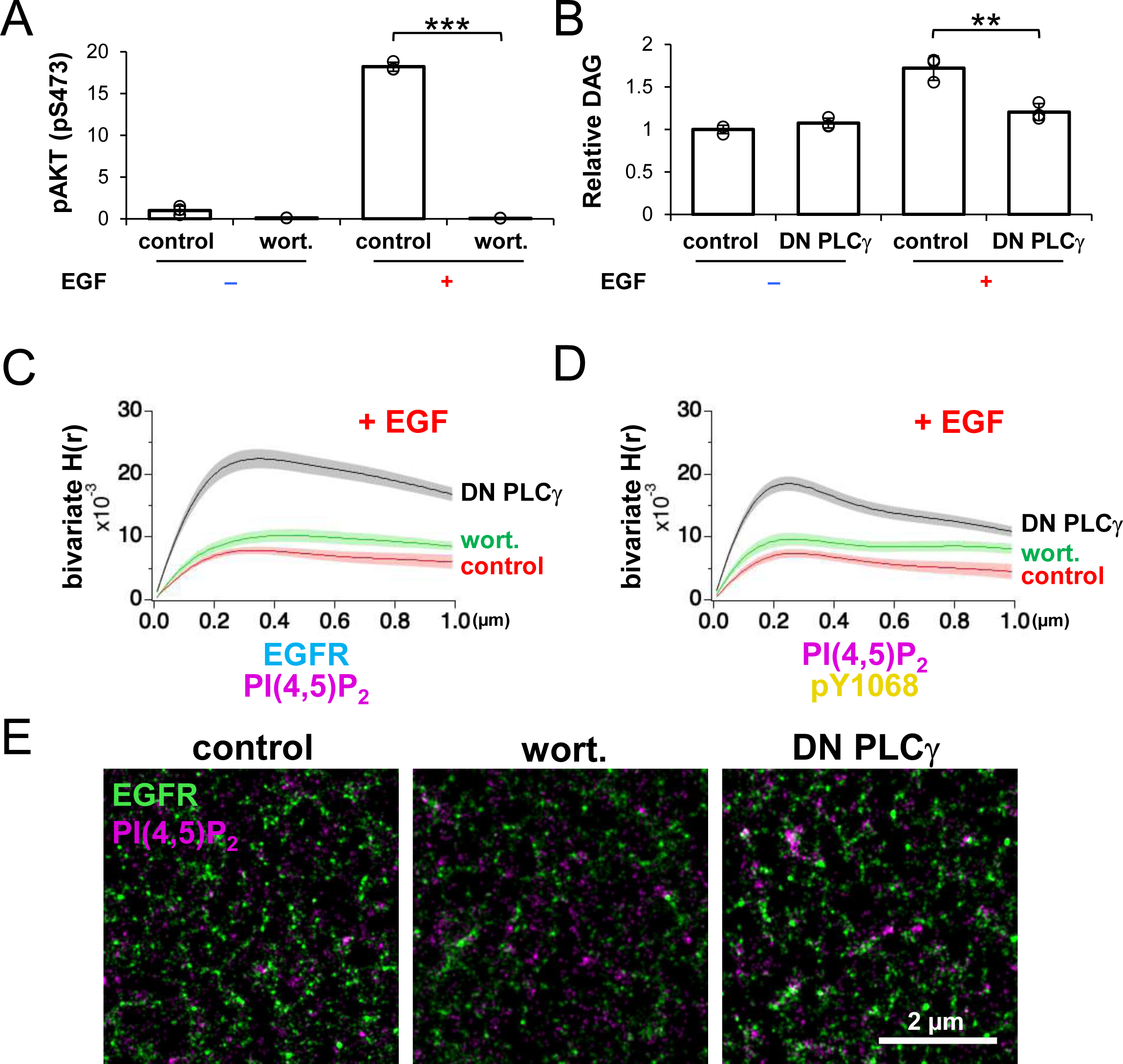
PLCγ is responsible for EGFR dissociation from PI(4,5)P_2_ nanodomains after EGFR is phosphorylated. (**A**) Effect of wortmannin on phosphorylation of AKT. A cell line stably expressing EGFR– rsKame was incubated in serum-free medium overnight and treated with 10 µM wortmannin for 1 h. The cells were then stimulated with or without 20 nM EGF for 1 min. Phosphorylated AKT was detected with an anti-phospho-AKT (Ser473) antibody in a western blotting analysis. Relative intensity was normalized to the mean value of the control cells without EGF stimulation. Data are means ± SD of three experiments. (**B**) Effect of DN PLCγ on DAG production. DN PLCγ was transiently expressed in a cell line stably expressing EGFR–rsKame. Cells were incubated in serum-free medium overnight and treated with or without 20 nM EGF for 1 min. Amounts of DAG were measured with a DAG assay kit. Relative intensity was normalized to the mean intensity of the control cells without EGF stimulation. Data are means ± SD of three experiments. (**C**) Bivariate H(r) value of EGFR–rsKame and PAmCherry–PI(4,5)P_2_ in control (red), wortmannin-treated (green), and DN-PLCγ-expressing cells (black) after incubation with 20 nM EGF for 1 min. In control and wortmannin-treated cells, PAmCherry1–PLCδ–PH was transiently expressed in a cell line stably expressing EGFR–rsKame. To co-express PAmCherry1–PLCδ–PH and DN PLCγ in the same cells, plasmid DEST40/PAmCherry1–PLCδ–PH–IRES–DN PLCγ was transfected to the cell line stably expressing EGFR–rsKame. Cells were incubated in serum- free medium overnight and treated with (control and DN PLCγ) or without 10 µM wortmannin for 1 h (wortmannin). The cells were then stimulated with 20 nM EGF for 1 min and treated with paraformaldehyde and glutaraldehyde. Data are means ± SEM of seven (control and wortmannin) or eight cells (DN PLCγ). (**D**) Bivariate H(r) value of PAmCherry–PI(4,5)P_2_ and HMSiR–pY1068 in control (red), wortmannin-treated (green), and DN-PLCγ-expressing cells (black) after incubation with 20 nM EGF for 1 min. After cells were prepared as described in (E), phosphorylated EGFR was immunostained with anti-EGFR (pY1068) and HMSiR-labeled antibodies. Data are means ± SEM of six (control), seven (wortmannin), or eight cells (DN PLCγ). (**E**) Images of PAmCherry–PI(4,5)P_2_ (magenta) and HMSiR-labeled pY1068 EGFR (green) after stimulation with EGF for 1 min. Cells were prepared as described in (F). **p < 0.01, ***p < 0.001 on Welch’s *t* test.

Under these conditions, we observed EGFR and PI(4,5)P_2_ in the plasma membrane with SMLM after EGF stimulation. To coexpress the PI(4,5)P_2_ probe and DN PLCγ in the same cells, we used a plasmid containing an internal ribosome entry site (IRES).

Measurement of Ripley’s bivariate H-function revealed that no significant difference in the bivariate H(R) value of EGFR and PI(4,5)P_2_ was observed between the wortmannin-treated (0.0102 ± 0.0010, n = 7 cells) and -untreated cells (0.0080 ± 0.0006, n = 7 cells), suggesting that PI3K is not important for the disaggregation of EGFR and PI(4,5)P_2_ (Fig. 6C). However, the high bivariate H(R) value of EGFR and PI(4,5)P_2_ was maintained in the DN-PLCγ-expressing cells (0.0225 ± 0.0015, n = 8 cells, p < 0.001) after EGF stimulation, suggesting that PLCγ is responsible for the reduced coaggregation of EGFR and PI(4,5)P_2_ (Fig. 6C).

We examined whether EGFR activation is truly associated with its coaggregation with PLCγ by observing phosphorylated EGFR and PI(4,5)P_2_ in the plasma membranes of wortmannin-treated or DN-PLCγ-expressing cells. After phosphorylated EGFR was immunostained with anti-EGFR (pY1068) and HMSiR-labeled antibodies in PAmCherry– PI(4,5)P_2_-expressing cells after EGF stimulation, the bivariate H(r) value of phospho-EGFR and PI(4,5)P_2_ was measured (Fig. 6D). The bivariate H(R) values of phospho-EGFR and PI(4,5)P_2_ in the control, wortmannin-treated, and DN-PLCγ-expressing cells were 0.0080 ± 0.0006 (n = 6 cells), 0.0093 ± 0.0010 (n = 7 cells), and 0.0193 ± 0.0012 (n = 8 cells), respectively. In the SMLM images of DN-PLCγ-expressing cells, overlapping of the nanodomains of phospho-EGFR and PI(4,5)P_2_ was observed in the submicrometer coaggregation domains indicated in the bivariate H-function analysis (Fig. 6E).

We found that the univariate H(R) values of PI(4,5)P_2_ were larger in the wortmannin-treated cells (0.071 ± 0.003, n = 9 cells) and DN-PLCγ-expressing cells (0.056 ± 0.005, n = 9 cells) than in the control cells (0.024 ± 0.002, n = 9; Fig. S7A). These results were expected from the inhibition of the metabolism of PI(4,5)P_2_ in the wortmannin-treated and DN-PLCγ-expressing cells (Fig. 6A and 6B). Interestingly, the R value was smaller in the wortmannin-treated cells (0.131 ± 0.013 µm, n = 9 cells) than in the DN-PLCγ-expressing cells (0.252 ± 0.005 µm, n = 9 cells, Fig. S7A), suggesting that PI3K and PLCγ affect the PI(4,5)P_2_ nanodomains in the plasma membrane differently. We detected no significant difference in the univariate H(R) values of pY1068 between the cells (Fig. S7B). These results support the conclusion that of the two enzymes PI3K and PLCγ, PLCγ mainly affects the reduction in the degree of coaggregation of EGFR and PI(4,5)P_2_ after EGFR phosphorylation.

ERBB3 directly activates PI3K through canonical p85 binding motifs in ERBB3 (Soltoff et al., 1994). One possible explanation for the different effects of these two enzymes is that PLCγ hydrolyzes PI(4,5)P_2_ around EGFR, whereas PI3K phosphorylates PI(4,5)P_2_ around the heterodimer of ERBB3 and EGFR in the plasma membrane. In EGFR KO and ERBB3 KO cells, little AKT phosphorylation was detected after EGF stimulation (Fig. S7C and S7D). EGFR–Halo expression restored AKT phosphorylation after EGF stimulation in EGFR KO cells, but not in ERBB3 KO cells. Therefore, both ERBB3 and EGFR were required for EGF-induced AKT phosphorylation under our experimental conditions. SMT analysis indicated that the colocalization rate of PLCγ–PI3K was lower than that of EGFR–PLCγ or EGFR–PI3K (Fig. S7E and S7F). These results support the idea that PLCγ and PI3K react differently to PI(4,5)P_2_ nanodomains in the plasma membrane.

### Hydrolysis of PI(4,5)P_2_ by PLC**_γ_** is involved in the deactivation of EGFR

As shown in Fig. 6, we found that PLCγ is responsible for the reduced coaggregation of EGFR and PI(4,5)P_2_ after EGF stimulation. PLCγ produces DAG, which is known to activate protein kinase C (PKC) (Steinberg, 2008). PKC activation is known to induce the phosphorylation of EGFR-Thr654 (Hunter et al., 1984), which is involved in the deactivation of EGFR. Therefore, we speculated that the hydrolysis of PI(4,5)P_2_ around EGFR molecules by PLCγ upon EGF stimulation has a role in the deactivation of EGFR via PKC. To test this hypothesis, we measured the amount of Thr654-phosphorylated EGFR under PLCγ-inhibited conditions (Fig. 7A and 7B). We found that EGFR Thr654 was less phosphorylated in DN-PLCγ-expressing cells than in the control cells after EGF stimulation. The amount of Thr654-phosphorylated EGFR was also reduced in PLCG1- knockdown (KD) cells. The reduced phosphorylation of EGFR-Thr654 in PLCG1-KD cells was restored by adding 4β-phorbol 12-myristate 13-acetate (PMA), which is a DAG mimetic (Fig. 7C and 7D), suggesting that the phosphorylation of Thr654 depends on DAG produced by PLCG1. Moreover, when the tyrosine kinase activity of EGFR was inhibited by AG1478, the phosphorylation of Thr654 was blocked in the control cells as well (Fig. 7C and 7D), which suggests that the phosphorylation of EGFR-Thr654 requires autophosphorylation of EGFR.

**Figure 7.**
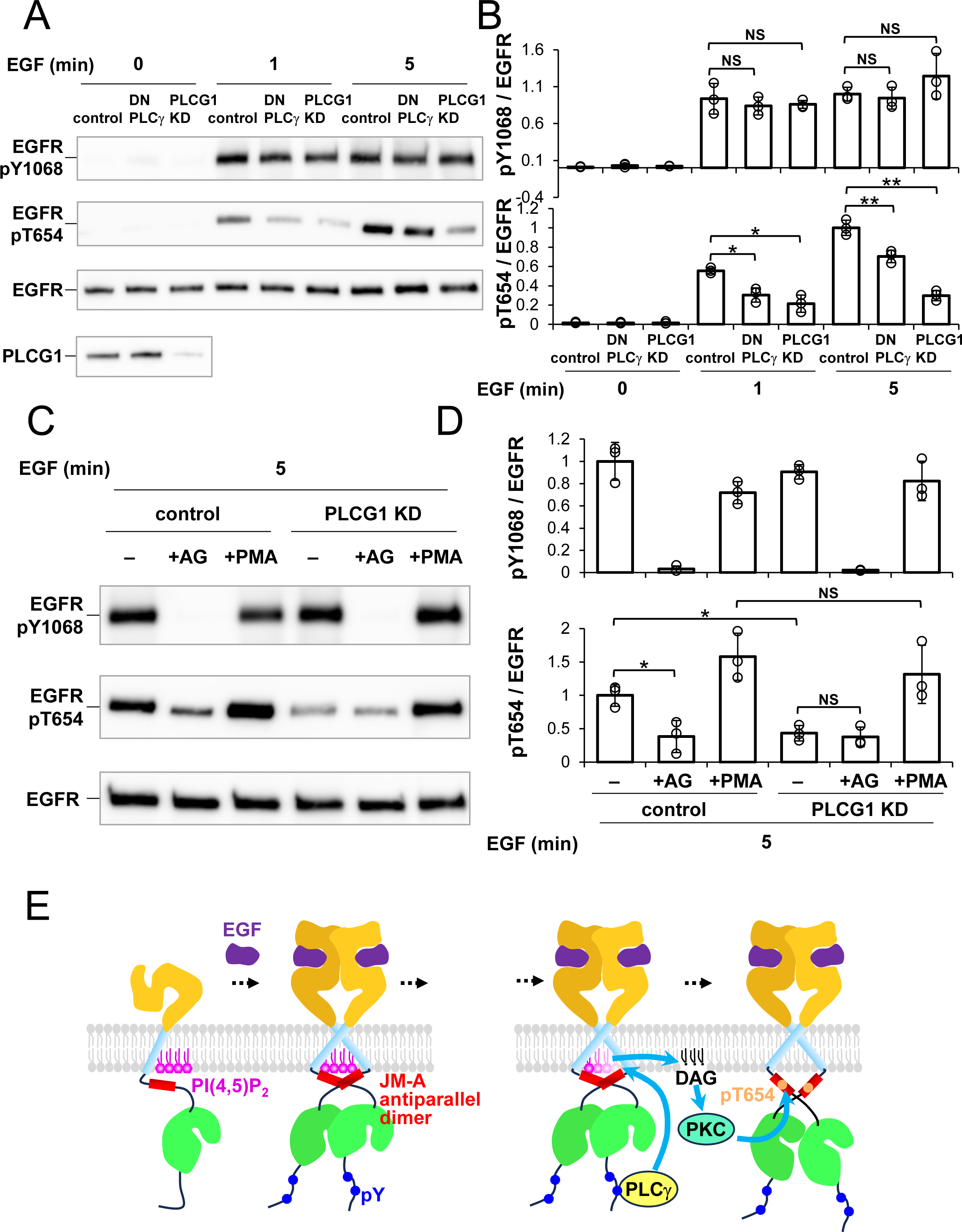
Thr654 of EGFR is phosphorylated in a PLCγ-dependent manner. (**A**) Effect of PLCγ on phosphorylation of EGFR. After EGFR-KO cells were transfected with EGFR–Halo, EGFR–Halo and DN PLCγ, or EGFR–Halo and PLCG1 siRNA, the cells were incubated in serum-free medium overnight and stimulated with 10 nM EGF for 1 min or 5 min. DN PLCγ was transiently expressed in EGFR-KO cells expressing EGFR–Halo. Phosphorylated-Tyr1068 EGFR, phosphorylated-Thr654 EGFR, and total EGFR were detected with anti-EGFR pY1068, anti-EGFR pT654, and anti-EGFR antibodies, respectively. (**B**) Ratios of phosphorylated-Tyr1068 EGFR/total EGFR and phosphorylated-Thr654 EGFR/total EGFR. The ratios were normalized to the mean value of lane #7 in (A). Data are means ± SD of three experiments. (**C**) Effects of AG1478 and PMA on phosphorylation of EGFR. After EGFR-KO cells were transfected with EGFR–Halo or EGFR–Halo and PLCG1 siRNA, the cells were incubated in serum-free medium overnight and stimulated with 10 nM EGF for 5 min in the presence or absence of 100 nM PMA. For inhibition of autophosphorylation of EGFR, cells were treated with 1 µM AG1478 for 30 min and stimulated with 10 nM EGF for 5 min. Phosphorylated-Tyr1068 EGFR, phosphorylated-Thr654 EGFR, and total EGFR were detected with anti-EGFR pY1068, anti-EGFR pT654, and anti-EGFR antibodies, respectively. (**D**) Ratios of phosphorylated-Tyr1068 EGFR/total EGFR and phosphorylated-Thr654 EGFR/total EGFR. The ratios were normalized to the mean value of lane #1 in (C). Data are means ± SD of three experiments. (**E**) Schematic model of the activation and signal transduction process for EGFR. Left, after EGF stimulation, PI(4,5)P_2_ (magenta) interacts with the arginine residues in the JM-A region (red) of EGFR, stabilizing the JM-A antiparallel dimer, and inducing the formation of asymmetric dimers of the kinase region (green) of EGFR. This dimerization results in the phosphorylation of several tyrosine residues in the tail region of EGFR. Right, Phosphorylated-Tyr1068 EGFR recruits PLCγ (yellow), degrading PI(4,5)P_2_ and producing DAG. DAG activates protein kinase C (cyan), which phosphorylates Thr654 of EGFR, deactivating EGFR. *p < 0.05, **p < 0.01, NS (not significant) on Welch’s *t* test.

To exclude the possibility that the inhibition of PLCγ reduces the autophosphorylation of EGFR, we measured the phosphorylation of EGFR-Tyr1068 in DN- PLCγ-expressing and PLCG1-KD cells (Fig. 7A and 7B). Consistent with previous literature on mouse fibroblasts (Ji et al., 1998), the levels of Tyr1068-phosphorylated EGFR were not significantly altered in the DN-PLCγ-expressing or PLCG1-KD cells, suggesting that the inhibition of PLCγ does not block the kinase activity of EGFR. These results suggest that the autophosphorylation of EGFR activates PLCγ after EGF stimulation, inducing the phosphorylation of EGFR-Thr654 in a DAG-dependent manner.

## Discussion

In this study, we examined the dynamics and functional roles of PI(4,5)P_2_ as the boundary and/or neighboring lipid of EGFR in the activation of EGFR by EGF stimulation. Superresolution microscopy revealed that EGFR and PI(4,5)P_2_ form nanodomains that partially overlap before EGF stimulation. The nanodomains also form domain clusters and the extent of coaggregation of EGFR and PI(4,5)P_2_ molecules in a higher-order domain structure decreases after EGF stimulation. Single-molecule imaging confirmed these results, detecting the transient colocalization of EGFR and PI(4,5)P_2_. The expression of a PI(4,5)P_2_ phosphatase reduced the dimerization and autophosphorylation of EGFR after EGF stimulation. These findings are consistent with previous *in vitro* and *in silico* observations that PI(4,5)P_2_ interacts strongly with the arginine residues in the JM-A region of EGFR, stabilizing the active conformation of EGFR (Abd Halim et al., 2015; Matsushita et al., 2013). Together with these observations, our biochemical analyses suggest that the local distribution of PI(4,5)P_2_ around EGFR before EGF stimulation not only acts as the substrate of EGFR effectors, including PI3K and PLCγ, after EGF stimulation, but also plays a crucial role in enhancing the dimerization and kinase activation of EGFR immediately after EGF stimulation (Fig. 7E, left).

We found that PI(4,5)P_2_ is also involved in a specific pathway of EGFR deactivation after EGF stimulation. Tyr-phosphorylated EGFR recruits PLCγ, degrading PI(4,5)P_2_ and producing DAG. DAG activates protein kinase C, which phosphorylates Thr654 of EGFR, deactivating EGFR (Fig. 7E, right). We have reported previously that Thr654 phosphorylation causes the JM-A dimer to dissociate in the presence of acidic lipids (Maeda et al., 2018; Maeda et al., 2022). This constitutes the negative feedback regulation of the EGFR kinase function. Consistent with this, in our experiment, the expression of DN PLCγ did not induce DAG production (Fig. 6B) or Thr654 phosphorylation (Fig. 7A and 7B), but increased the Tyr1068 phosphorylation of EGFR (Fig. 7A and 7B). After the degradation of PI(4,5)P_2_, EGFR is free from both positive and negative regulation by PI(4,5)P_2_. We expect this state to facilitate the higher-order oligomerization of EGFR, which is important for signaling to cytoplasmic proteins (Hiroshima et al., 2018; Maeda et al., 2022).

We found that of PI3K and PLCγ, PLCγ is responsible for the disaggregation of the EGFR and PI(4,5)P_2_ nanodomains after EGFR phosphorylation. The difference between the effects of PI3K and PLCγ on the PI(4,5)P_2_ nanodomains can be explained as follows.

PLCγ is activated by EGFR after EGF stimulation (Carpenter and Ji, 1999), whereas PI3K is activated by ERBB3, which is associated with EGFR stimulation (Hellyer et al., 1998; Mattoon et al., 2004). This is supported by our results, which show that both ERBB3 and EGFR are required for EGF-induced AKT phosphorylation (Fig. S7C and S7D). Therefore, the EGFR–EGFR–PLCγ and EGFR–ERBB3–PI3K complexes exist separately in the plasma membrane after EGF stimulation, which is consistent with the results in Fig. S7E and S7F. Upon EGF stimulation, EGFR preferentially phosphorylates EGFR rather than ERBB3 (Okada et al., 2022). This may lead to higher activation of the EGFR–EGFR–PLCγ complex than the EGFR–ERBB3–PI3K complex. The small aggregation of PI(4,5)P_2_ nanodomains in wortmannin-treated cells (Fig. S7A) may indicate newly synthesized (An et al., 2022) or partially degraded PI(4,5)P_2_ nanodomains at the plasma membrane after EGF stimulation.

EGF stimulation does not induce a significant change in PI(4,5)P_2_ levels in the plasma membrane (Delos Santos et al., 2017). Here, we show that EGF stimulation reduces the density of small PI(4,5)P_2_ nanodomains in the plasma membrane around the EGFR. Together, these results indicate that PI(4,5)P_2_ is hydrolyzed by PLCγ only in the region surrounding EGFR. In contrast, synaptojanin expression reduces PI(4,5)P_2_ levels by 45% (Field et al., 2005), leading to defects in cytokinesis and mislocalization of PI(4,5)P_2_- binding proteins to the plasma membrane (Abe et al., 2012; Abe et al., 2021; Field et al., 2005). These results suggest that synaptojanin expression decreases the amount of PI(4,5)P_2_ throughout the plasma membrane rather than in the restricted region around EGFR.

We have shown here that the degree of coaggregation between EGFR and PI(4,5)P_2_ is higher than that between EGFR and PS before EGF stimulation. The preferential coaggregation of the EGFR nanodomains with PI(4,5)P_2_ rather than PS is probably due to the following two factors. The first is the curvature of the membrane. Previous electron microscopy–bivariate analyses showed that the nanodomains of inactive HRAS were enriched with PI(4,5)P_2_, whereas those of active KRAS4B were enriched with PS due to highly specific RAS–lipid interactions determined by the membrane curvature (Liang et al., 2019; Zhou et al., 2014; Zhou et al., 2015). Like inactive HRAS, EGFR may prefer the PI(4,5)P_2_ nanodomain environment, in addition to interacting electrostatically with PI(4,5)P_2_. The second factor is tetraspanins, which form nanodomains at the plasma membrane. Tetraspanins may interact with EGFR and regulate EGFR activities and dynamics within tetraspanin nanodomains (Berditchevski and Odintsova, 2016; Odintsova et al., 2003; Sugiyama et al., 2023). Tetraspanins also indirectly interact with PI(4,5)P_2_ (Halova and Draber, 2016). Therefore, EGFR may be localized in PI(4,5)P_2_-enriched nanodomains mediated by tetraspanins. Further studies are required to reveal the relationships between these molecules.

PI(4,5)P_2_ nanodomains may suppress EGFR signaling in the resting state despite their stabilizing effect on the JM-A dimer of EGFR during its kinase activation. Several reports have suggested that membrane cholesterol prevents EGFR phosphorylation before EGF stimulation (Chen and Resh, 2002; Ringerike et al., 2002). A biochemical analysis suggested that EGFR is associated with the detergent-resistant membrane fraction, which is enriched with cholesterol and PI(4,5)P_2_ (Brown and London, 1998; Hope and Pike, 1996; Laux et al., 2000). We previously showed with SMLM that PI(4,5)P_2_ nanodomains appear in the inner leaflet just beneath the sphingomyelin/cholesterol nanodomains located in the outer leaflet of the plasma membrane (Abe et al., 2012; Makino et al., 2017). Together with the results of the present study, this suggests that cholesterol- and PI(4,5)P_2_-rich membrane domains exist in the plasma membrane, form higher-order aggregates, and transiently colocalize with EGFR clusters before EGF stimulation. Such membrane structures may prevent EGFR from activation because cholesterol exerts an inhibitory effect on EGFR. However, after EGF stimulation, rapid EGFR activation is induced by the positive effect of PI(4,5)P_2_ on the dimerization of EGFR kinase. Transient complex of EGFR dimers with PI(4,5)P_2_ may be required to leave the cholesterol nanodomains to allow kinase activation. The transient mobilization of EGFR clusters immediately after EGF stimulation that we have observed previously (Hiroshima et al. 2018) may reflect this process.

SMT analysis revealed that EGFR monomers decreased and dimers increased in the immobile fraction of control CHO-K1 cells after EGF stimulation (Fig. 4A). The changes in oligomer size were smaller in the slow- and fast-mobile fractions than in the immobile fraction. In addition, the fraction size of the immobile state was not reduced after EGF stimulation (Fig. 3B). These results suggest that stable EGFR dimer/oligomers were mostly increased in the immobile fraction after EGF stimulation. In contrast, a decrease in the colocalization rate between EGFR and PI(4,5)P_2_ after EGF stimulation was observed in the slow- and fast-mobile fractions but not in the immobile fraction (Fig. S3D). The PI(4,5)P_2_ probes employed in this study detect PI(4,5)P_2_ near EGFR but they might not bind to PI(4,5)P_2_ associated with EGFR due to steric hindrance. It is plausible that concentrated PI(4,5)P_2_ molecules help to stabilize EGFR dimer/oligomers in the immobile fraction. However, we cannot exclude the possibility that the dimer/oligomers in the slow- and fast-mobile fractions were also stabilized by PI(4,5)P_2_, which was not detected by the PI(4,5)P_2_ probes.

For SMT analysis, we used EGFR–GFP rather than antibody-mediated labeling, as in previous studies (Mudumbi et al., 2024; Sugiyama et al., 2023). Compared with antibody-mediated labeling in the extracellular region of EGFR (Sugiyama et al., 2023), a higher change in EGFR mobility upon EGF stimulation was observed. The characteristics of EGFR or its binding efficiency to EGF may differ between the labeling methods.

Needham et al. (2016) and Mudumbi et al. (2024) reported an increase in the size of single EGFR clusters after EGF stimulation as an indication of clustering using superresolution microscopy (Mudumbi et al., 2024; Needham et al., 2016). Such faint increases were difficult to observe in our SMLM imaging under dense EGFR expression (Fig. 1); however, SMT analysis detected increases in the singe-spot fluorescence intensities (Fig. 4) (Hiroshima et al., 2018; Yasui et al., 2018), which is consistent with EGF-induced EGFR clustering.

In conclusion, PI(4,5)P_2_ plays a crucial role as a molecular switch for EGFR functions. PI(4,5)P_2_ is involved in the complicated remodeling of the boundary lipids and higher-order membrane structure around EGFR. To detect whole these processes directly, further observations are required using cholesterol and DAG probes under SMLM or electron microscopy. Alternatively, further biochemical analyses measuring the local lipid contents around EGFR should be performed.

## Materials and Methods

### Cell Culture and Drug Treatments

HeLa cells were grown at 37°C in Dulbecco’s modified Eagle’s medium (DMEM) (Nacalai Tesque Inc., Kyoto, Japan) supplemented with 10% fetal bovine serum (FBS).

CHO-K1 cells were grown at 37°C in Ham’s F-12 medium (FUJIFILM Wako Pure Chemical Corporation, Osaka, Japan), supplemented with 10% FBS. All EGF stimulations were performed at 25°C after preincubation of the cells at 25°C for at least 10 min.

### Plasmid construction

DEST40/rsKame was generated by introducing V157L mutation into Dronpa in DEST40/Dronpa (Abe et al., 2012). The coding sequence for human EGFR was cloned into DEST40/rsKame to generating DEST40/EGFR–rsKame. The coding sequence for the human GRP1-PH region was obtained from HeLa cell cDNA by PCR amplification and cloned into DEST40/PAmCherry1 to generate DEST40/PAmCherry–GRP1-PH. The coding sequence for the z-region of human PLCγ (DN PLCγ) was cloned into DEST40 (Abe et al., 2012) to generate DEST40/DN PLCγ. To generate DEST40/PAmCherry– PLCδ–PH–IRES–DN PLCγ, fragments of IRES and DN PLCγ were introduced immediately after PLCδ–PH in DEST40/PAmCherry– PLCδ–PH (Abe et al., 2012). Halo– PLCδ–PH, Halo–evt–2–PH, and Halo–TubbyC were generated by replacing the fragment of GRB2 in Halo–GRB2 (Yasui et al., 2018) with PLCδ–PH, evt–2–PH, and TubbyC, respectively. EGFR–Halo was generated by replacing the GFP fragment in EGFR–GFP (Hiroshima et al., 2018) with Halo7. EGFR(3RN)–Halo was generated by introducing R645N/R646N/R647N mutations into EGFR in EGFR–Halo. EGFRvIII–Halo was generated by deleting 30-297 amino acids of EGFR in EGFR–Halo. The coding sequence for human PLCG1 was obtained from HeLa cell cDNA by PCR amplification. PLCG1– SNAP was generated by replacing the EGFR and halo fragments in EGFR–Halo with PLCG1 and SNAP, respectively.

### Generation of EGFR or ERBB3-KO cells with CRISPR/Cas9 gene editing

To construct the gene-editing plasmids, oligomers (5′- CACCGCACAGTGGAGCGAATTCCTT-3′ and 5′- AAACAAGGAATTCGCTCCACTGTGC-3′ for EGFR, 5′- CACCGTGATCCAGCAGAGAACCCAG-3′ and 5′-AAACCTGGGTTCTCTGCTGGATCAC-3′ for ERBB3) were synthesized, annealed, and cloned into the PX459 vector (#48139, Addgene) at the BbsI site, as previously described (Ran et al., 2013). HeLa cells were transfected with the resultant plasmid using Lipofectamine 3000 (Thermo Fisher Scientific, Waltham, MA). Single colonies were picked after selection with 0.3 μg/ml puromycin for three days. The genomic DNAs were extracted from the clones with the GenElute™ Mammalian Genomic DNA Miniprep Kit (Sigma-Aldrich, St. Louis, MO). The genome sequences were examined by PCR using the following primers: 5′-GATCGTGGACATGCTGCCTCCTGTGTCCATGACTGC-3′ and 5′-CTTCCCCTGCAGTATCTTACACACAGCCGGC-3′ for EGFR KO, or 5′- AGGTGGGGAAGGCATCTAGGGCAAAGGG-3′ and 5′- CGGAACTCGGGCGGTAATGCAAGTGATGG-3′ for ERBB3 KO cells. Sequence analysis revealed that EGFR KO had a 1-bp deletion (C) next to the protospacer adjacent motif (PAM) sequence and that ERBB3 KO had a 1-bp deletion (C) next to the PAM sequence.

### Establishing cell lines stably expressing EGFR–rsKame

To establish cell lines stably expressing EGFR–rsKame, EGFR KO cells were transfected with DEST40/EGFR–rsKame using Lipofectamine 3000 (Thermo Fisher Scientific) and cultured in the presence of 1 mg/ml G418 (Nacalai Tesque Inc.) for 14 days. Stable clones expressing EGFR–rsKame were selected using fluorescence microscopy.

### Western blotting analysis

Parental HeLa and EGFR-KO cells stably expressing the EGFR–rsKame fusion protein were incubated in DMEM supplemented with 10% FBS. EGFR-KO cells were transfected with EGFR–GFP and incubated in DMEM supplemented with 10% FBS for 24 h. EGFR-KO and ERBB3-KO cells were transfected with EGFR–Halo and incubated in DMEM supplemented with 10% FBS for 24 h. The cells were further incubated in serum-free medium overnight and stimulated with 20 nM EGF for 1 min. After the cells were lysed in ice-cold RIPA lysis buffer (Nacalai Tesque Inc.), the lysates were cleared by centrifugation at 15,000 × g for 10 min at 4°C. Protein concentrations were determined using a Pierce™ BCA Protein Assay Kit (Thermo Fisher Scientific). Proteins were applied on 4–15% TGX precast gels (Bio-Rad Laboratories, Hercules, CA) so that the protein concentrations were the same. The gels were transferred onto PVDF membranes using the Trans-Blot Turbo Transfer System and blocked with 2% non-fat dry milk in TBS + 0.1% Tween (TBST). Total EGFR was detected using anti-EGFR (#sc-3; Santa Cruz

Biotechnology; 1:500) as the primary antibody and horseradish peroxidase (HRP)-linked anti-rabbit IgG (#7074, Cell Signaling Technology, Danvers, MA; 1:1000) as the secondary antibody. Phosphorylated EGFR (pY1068) was detected using anti-EGFR pY1068 (#3777, Cell Signaling Technology; 1:1000) as the primary antibody and HRP-linked anti-rabbit IgG. Total EGFR was detected as described above. Phosphorylated EGFR (pY1173) was detected using anti-EGFR pY1173 (sc-12351; Santa Cruz Biotechnology, Dallas, TX, USA; 1:500) as the primary antibody and HRP-linked anti-goat IgG (sc-2354, Santa Cruz Biotechnology; 1:1000) as a secondary antibody. Phosphorylated AKT (pS473) was detected using anti-AKT (S473) (#4060, Cell Signaling Technology; 1:1000) as the primary antibody and HRP-linked anti-rabbit IgG. Phosphorylated ERK (Thr202/Tyr204) was detected using anti-phospho-p44/42 (#4370, Cell Signaling Technology; 1:1000) as a primary antibody and HRP-linked anti-rabbit IgG. Total ERK was detected using anti- p44/42 (#4695, Cell Signaling Technology; 1:1000) as the primary antibody and HRP-linked anti-rabbit IgG. Immunoreactive proteins were detected with ECL Prime Western Blotting Detection Reagent (GE Healthcare) using an ImageQuant LAS 500 device (GE Healthcare).

### Analysis of the effects of PLC**γ** and PKC on EGFR phosphorylation

EGFR-KO cells were transfected with EGFR(WT)–Halo, GFP–SYNJ, EGFR(3RN)–Halo, DN PLCγ, or PLCG1 siRNA (MISSION® siRNA, SASI_Hs02_00334210, Sigma-Aldrich) using the Neon Transfection System (Thermo Fisher Scientific) and incubated in DMEM supplemented with 10% FBS for 24 h. The medium was replaced with DMEM supplemented with 0.2% bovine serum albumin (BSA), and the cells were further incubated for 24 h and stimulated with EGF. For PMA treatment, the cells were stimulated with EGF in the presence of 100 nM PMA. To inhibit autophosphorylation of EGFR, cells were pretreated with 1 µM AG1478 for 30 min and stimulated with EGF. Cells were washed twice with cold PBS. After the cells were lysed as described above, the proteins were separated on 4–15% TGX precast gels. Total EGFR was detected as described above. After the intensity of total EGFR was measured, we applied the proteins to SDS-PAGE again, so that the intensity of total EGFR in each sample was the same. Phosphorylated EGFR (pT654) was detected using anti-EGFR pT654 (#ab75986; Abcam, Cambridge, UK; 1:500) as the primary antibody and HRP-linked anti-rabbit IgG. PLCG1 was detected using anti-PLCγ1 (#5690, Cell Signaling Technology; 1:500) as the primary antibody and HRP-linked anti-rabbit IgG. Phosphorylated EGFR (pY1068) and total EGFR were detected, as described above.

### Measurement of DAG

DN PLCγ was transiently expressed in a cell line stably expressing EGFR– rsKame. The cells were incubated in serum-free medium overnight and treated with or without 20 nM EGF for 1 min. The amount of DAG in the DN PLCγ-expressing and control cells was measured using a DAG assay kit (#MET-5028, Cell Biolabs, Inc. San Diego, CA) according to the manufacturer’s instructions.

### Inhibition of PI3K

A cell line stably expressing EGFR–rsKame was incubated in serum-free medium overnight and treated with or without 10 µM wortmannin for 1 h. The cells were stimulated with or without 20 nM EGF for 1 min. Phosphorylated AKT was detected by western blotting analysis as described above. For SMLM imaging, after EGF stimulation, cells were treated with paraformaldehyde and glutaraldehyde.

### Analysis of EGFR dimerization

After EGFR KO cells were transfected with EGFR(WT)–Halo or EGFR(3RN)– Halo, the cells were incubated in DMEM supplemented with 10% FBS for 24 h. The medium was replaced with DMEM supplemented with 0.2% BSA, and the cells were further incubated for 24 h. The cells were incubated with 0.2 or 20 nM EGF for 2 min and washed twice with cold PBS. After the cells were incubated with 3 mM BS3 (Thermo Fisher Scientific) in PBS at 4°C for 1h, BS3 was quenched with 20 mM Tris-HCl (pH 8.0). The cells were then washed twice with cold phosphate-buffered saline (PBS). After the cells were lysed, protein concentrations were determined. Proteins were subjected to western blotting analysis so that the protein concentrations were the same.

### Cy5–EGF labeling

After EGFR-KO cells were transfected with EGFR(WT)–Halo, EGFR(3RN)– Halo, or EGFRvIII–Halo, cells were incubated in DMEM supplemented with 10% FBS for 24 h. The medium was replaced with DMEM supplemented with 0.2% BSA, and cells were further incubated for 24 h. Cells were stained with 10 nM Janelia Fluor® 549 (JF549) HaloTag Ligand (Promega, Madison, WI) for 30 min and were washed with the medium at least three times. After the medium was replaced with DMEM/Ham’s F-12 containing HEPES and 0.2% BSA, 10 ng/mL Cy5–EGF was added as described previously (Sako et al., 2000). Confocal images were obtained within 10 min after Cy5–EGF with a confocal microscope (FV 3000, Olympus, Tokyo, Japan) equipped with a 60 × 1.35 NA objective lens.

### Single-molecule tracking (SMT) analysis

SMT analysis was performed as previously described (Kuwashima et al., 2021; Yanagawa et al., 2018; Yanagawa and Sako, 2021) with modifications. For SMT analysis of EGFR and PI(4,5)P_2_, HeLa cells were transfected with EGFR–GFP and Halo-PLCδ–PH. After the cells were incubated in DMEM supplemented with 10% FBS for 24 h, the medium was replaced with DMEM supplemented with 0.2% BSA and the cells were further incubated for 24 h. Cells were stained with 10 nM JF549 for 30 min and washed with DMEM/Ham’s F-12 containing HEPES and 0.2% BSA, at least three times. Single-molecule imaging of the living cells was performed at 25 °C. Fluorescently labeled proteins were observed under a microscope (TiE, Nikon, Tokyo, Japan). The cells were illuminated with a 488-nm laser (OBIS 488, Coherent Inc., Santa Clara, CA, USA) and a 561-nm laser (OBIS 561, Coherent). Fluorescent images of 150-200 frames were recorded using an Auto Imaging System (Zido, Osaka, Japan; http://zido.co.jp/en/) with an exposure time of 30 ms for dual-color imaging. The SMT and Variational Bayesian-Hidden Markov Model clustering analyses were performed with the Auto Analysis System (Zido) based on a two-dimensional Gaussian fitting algorithm, as described previously (Kuwashima et al., 2021; Yanagawa et al., 2018; Yanagawa and Sako, 2021). All subsequent analyses (diffusion dynamics, intensity distribution, colocalization, and statistical analysis) were performed using an updated version of the smDynamicsAnalyzer, Igor Pro 9.0, and a WaveMetrix (Igor)-based homemade program. Refer to the reference for detailed instructions and curve fitting functions (Yanagawa and Sako, 2021). Colocalization analysis was performed with slight modifications to account for the differences in expression levels among cells. After measuring the density of the particles (i.e., particle number/cell area), we normalized the original colocalization rate (Yanagawa and Sako, 2021) to the densities. We normalized all data for the colocalization rate and presented them as relative colocalization rate.

For SMT analysis of EGFR under PI(4,5)P_2_-interaction-depleted conditions, CHO- K1 cells were transfected with EGFR–Halo, EGFR–Halo and DEST40/GFP–SYNJ (Abe et al., 2021), or EGFR(3RN)–Halo and incubated in Ham’s F-12 supplemented with 10% FBS for 24 h. After replacing the medium with Ham’s F-12 supplemented with 0.2% BSA, the cells were further incubated for 24 h. To observe EGFR in HeLa cells, EGFR-KO HeLa cells were transfected with EGFR–Halo or EGFR–Halo and DEST40/GFP–SYNJ and incubated in DMEM supplemented with 10% FBS for 24 h. After the medium was replaced with DMEM supplemented with 0.2% BSA, cells were further incubated for 24 h. Cells were stained with 1 nM SaraFluor™ 650T (SF650) HaloTag Ligand (#A308-01, GORYO Chemical, Inc. Sapporo, Japan) for 30 min and washed at least three times with DMEM/Ham’s F-12 containing HEPES and 0.2% BSA. The fluorescently labeled proteins were observed using a 637-nm laser (OBIS 637, Coherent Inc.). Expression of GFP–SYNJ was confirmed using a 488-nm laser. The fluorescence images were recorded as described above.

For SMT analysis of EGFR, PLCγ, and PI3K, EGFR-KO HeLa cells were transfected with EGFR–GFP, PLCG1–SNAP, and Halo–p85α and incubated in DMEM supplemented with 10% FBS for 24 h. After the medium was replaced with DMEM supplemented with 0.2% BSA, cells were further incubated for 24 h. Cells were stained with 10 nM SNAP-Cell TMR-Star ligand (#S9105S, New England Biolabs) and 1 nM SF650 HaloTag ligand for 30 min, and washed at least three times with DMEM/Ham’s F- 12 containing HEPES and 0.2% BSA. The fluorescently labeled proteins were observed under a microscope as described above with a 637-nm laser (OBIS 637, Coherent Inc.). The fluorescently labeled proteins were observed with 488-nm laser, 561-nm laser, and 637-nm laser. The fluorescent images were recorded as described above.

### SMLM imaging

Cell lines stably expressing EGFR–rsKame were transfected with the plasmids using the Neon Transfection System (Thermo Fisher Scientific). After 1 d, the cells were incubated with 10 nM HMSiR HaloTag Ligand (#A201-01, GORYO Chemical, Inc.) in DMEM supplemented with 0.2% BSA for 16 h. The cells were then washed with fresh medium and incubated for 3 h. The cells were fixed with 4% paraformaldehyde and 0.2% glutaraldehyde for 60 min and washed with PBS. To calibrate the drift, TetraSpeck™ Microspheres (#T7279; Thermo Fisher Scientific) were added before observation. Similar to the photoactivation of Dronpa, we photoactivated EGFR–rsKame on the basal plasma membrane with a 488-nm laser (OBIS 488, Coherent) for excitation and turning off the fluorescence, and attributed this to the spontaneous recovery of the stochastic turning-on of the fluorescence, instead of the illumination at 405 nm (Mizuno et al., 2010). PAmCherry was photoactivated with a 405-nm (COMPASS 405-25 CW, Coherent) and illuminated with a 532-nm laser (COMPASS 315M-100, Coherent). HMSiR was illuminated with a 637-nm laser (OBIS 637, Coherent). EGFR–rsKame, PAmCherry, and HMSiR were observed under total internal reflection illumination using an inverted fluorescence microscope (TiE). 1000–2000 frames of fluorescent images were acquired using the Auto Imaging System (Zido, Japan) at a frame rate of 30 ms.

To stain phosphorylated EGFR, cells were fixed with 4% paraformaldehyde and 0.2% glutaraldehyde for 60 min. Cells were permeabilized with 0.1% Triton X-100 in PBS and blocked with 2% BSA for 1 h. The cells were then incubated with anti-EGFR pY1068 (#3777, Cell Signaling Technology, 1:100) for 1 h. The cells were then washed and incubated with HMSiR-coupled goat anti-rabbit IgG (#A204-01, GORYO Chemical, Inc.).

### SMLM analysis workflow

All subsequent analyses (image reconstitution, automated multi-distance spatial clustering analysis, G-SMAP analysis, and statistical analysis) were performed using an updated version of smDynamicsAnalyzer. See the legend for Fig. 1S and Yanagawa and Sako, 2021 for the details of SMLM analysis workflow.

Localizations of single-fluorophore-labeled proteins (samples) and TetraSpeck™ Microspheres (beads) were determined by 2D Gaussian fitting algorism. Thermal drift and misalignment between color channels were corrected by an affine transformation using the localizations of the beads as indices (Fig. S1 B). The positional accuracy of the immobile fluorescence beads was estimated to be 6-8 nm (full width at half maximum, FWHM) (Figs. S1B and S2D). The positional accuracy of the single molecules was typically 20-30 nm. The accuracy of the superimposition of two images after the affine transformation was 10-14 nm (Figs. S1B and S2E).

SMLM images were reconstituted by convolving 2D-Gaussian functions for the single-molecule images (Fig. S1C). To avoid multi-counting of the same molecule, the positions of a single-molecule were averaged and plotted at a single location. The image resolution of SMLM was estimated with a method based on Fourier ring correlation (FRC) (Nieuwenhuizen et al., 2013). Resolution values for Dronpa(V157L), PAmCherry1, and HMSiR were 30.1 ± 2.9 nm, 26.8 ± 1.2 nm, and 28.2 ± 1.0 nm, respectively (means ± SEM, n = 5).

We developed and used a multi-distance spatial cluster analysis program to estimate the size and distribution of nano-domains automatically (Fig. S1D-S1F), where the local point density around each point was compared with that for the random distribution based on the Ripley’s K-function, K(r), and its variants for cluster analysis (Kiskowski et al., 2009) and co-cluster analysis (Lagache et al., 2018). We also generated a G-function spatial map (G-SMAP) to visualize single clusters (Fig. S1 F). The size of each cluster, the number of points within each cluster, and the point density in a cluster were calculated in the G-SMAPs.

### Statistical analysis

Statistical analyses were performed using Igor Pro software (WaveMetrics, Inc., Lake Oswego, OR) or Microsoft Excel (Microsoft, Redmond, WA). The experiments were repeated at least three times, as described in the figures and corresponding figure legends. The results are expressed as the means ± SEM. Statistical significance was determined using Tukey’s multiple comparison test or Welch’s t-test between the groups. Statistical significance was set at p < 0.05, and they were classified into the following four categories: * (p < 0.05), ** (p < 0.01), *** (p < 0.001), and NS (not significant, p ≥ 0.05).

## Supplemental materials

Fig. S1 shows the multicolor SMLM analysis workflow. Fig. S2 shows the construction and evaluation of multicolor SMLM. Fig. S3 shows SMT analysis of EGFR– GFP and JF549–PI(4,5)P_2_. Fig. S4 shows time-dependent colocalization of EGFR-PI(4,5)P_2_ and EGFR-GRB2. Fig. S5 shows SMT analysis of EGFR under PI(4,5)P_2_-depleted condition. Fig. S6 shows SMT analysis of EGFR under PI(4,5)P_2_-interaction-depleted condition. Fig. S7 shows that PI3K is not responsible for EGFR dissociation from PI(4,5)P_2_ nanodomains after EGFR is phosphorylated.

Movie1 shows the lateral colization of EGFR and PI(4,5)P_2_ decreases after EGF stimulation. EGFR–GFP (green) and Halo–PI(4,5)P_2_ (magenta) were transiently expressed in EGFR-KO HeLa cells. After the cells were incubated in serum-free medium overnight, Halo–PI(4,5)P_2_ was stained with JF549–Halo ligand. EGFR–GFP and JF549–PI(4,5)P_2_ were observed at a time resolution of 30 ms for 6 s before EGF stimulation (left). After stimulation with 20 nM EGF, EGFR–GFP and JF549–PI(4,5)P_2_ were observed in the same cell 2 min after the addition of EGF (right).

## Data availability

The raw data and code are available from the corresponding author upon reasonable request.

## Acknowledgments

This work was supported by Grants-in-Aid from the Ministry of Education, Culture, Sports, Science, and Technology of Japan (22K06609 to M.A, 19H05647 to Y.S.); Japan Science and Technology Agency (JST), PRESTO, JPMJPR20EF (M,Y); and the Glycolipidologue Program of RIKEN (to Y.S. and T.K). We thank Dr. Ryo Maeda for the technical advice for biological experiments. We also thank Maiko Minatohara for biochemical experiments. We thank the Support Unit for Bio-Material Analysis, RRD, RIKEN for DNA sequence and cell sorting.

## Author contributions

M.A., M.Y., and Y.S. designed the experiments. M.H., M.Y., and Y.S. developed the microscope station. M.Y. developed the software for SMLM and SMT.

M.A. performed the biological experiments. M.A. and T.K. designed and made the lipid probes. M.A. and M.Y. performed SMLM and SMT experiments and data analysis. M.A., M.Y., and Y.S. wrote the manuscript with input from all co-authors.

---

**Figure S1.**
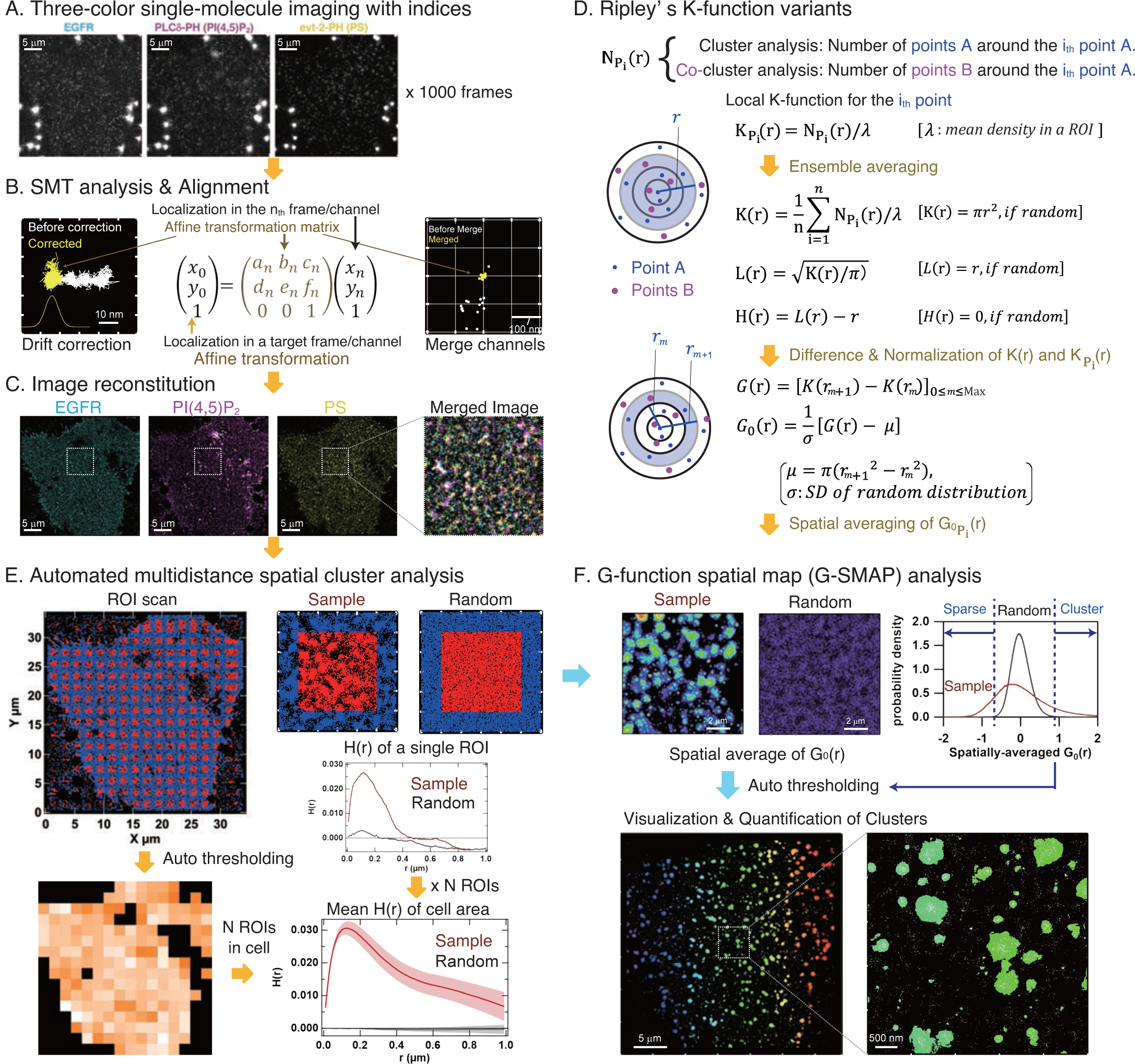
Multicolor SMLM analysis workflow. (**A**) Typical images of 512 × 512 pixels (34.3 × 34.3 µm with a pixel size of 67 nm). Bright spots around the cells indicate 100 nm TetraSpeck Microspheres (beads), and those in the cells indicate fluorescence-labeled EGFR, PI(4,5)P_2_, or PS (samples). (**B**) SMT analysis followed by alignment using affine transformation. An affine transformation matrix was calculated from the XY coordinates of the same beads in the n_th_ frame (or color channel) and a target frame (or channel) using least-squares regression. Example trajectories of the same beads before (white) and after (yellow) drift correction during 1000 frames are shown in the left panel. Histogram of the X coordinates of the beads after affine transformation is shown under the trajectory. The difference in the XY coordinates of the same beads between two color channels, before (white) and after (yellow) affine transformation, is shown in the right panel. (**C**) Reconstituted images. After the alignment of localization with affine transformation, images were reconstituted by convolution with a Gaussian kernel around each localization of the sample. The 5 × 5 µm area is merged and expanded in the right panel. (**D**) Definition and calculation flow of Ripley’s K-function variants. Np_i_(r) is the number of points within distance r around the i_th_ point. In the calculation of the univariate K-function, the number of points belonging to the same channel was counted. In the bivariate K-function calculation, the number of points belonging to the other channel was counted. Kp_i_(r) is a local K-function around the i_th_ point, where λ is the mean density in the region of interest (ROI). K(r) is Ripley’s K-function, which is the ensemble average of all Kp_i_(r) for n points in the ROI. K(r) is πr^2^ if the spatial distribution of the n points in the ROI is random. Because intuitive comparison with πr^2^ is difficult, variants L(r) and H(r) are often used; L(r) and H(r) become r and 0 if random, respectively. G(r) is the difference function of K(r), where the number of points in the donut area between circles of radii r_m+1_ and r_m_ is centered at each point. G_0_(r) is normalized G(r), where μ and σ are the mean and SD of G(r) in a random distribution. Unlike H(r), G(r) does not involve nonlinear computations, and therefore is suitable for the following spatial mapping analysis. (**E**) Automated multidistance spatial cluster analysis. After alignment with affine transformation, the image area was divided into ROIs with two rectangles (top panels). K(r) and its variants for the sample and for random simulation data were computed for the center points within the red region. In the calculation, the target points within both the red and blue areas were considered, to reduce the edge effect. After the calculations for all ROIs, ROIs containing cellular regions were detected with tiling images of the mean density of each ROI (lower left panel). The Otsu method was used to binarize the cellular and noncellular regions. The lower right panel shows an example of the mean H(r) for the cellular region. Data are shown as means ± SEM of N ROIs within the cellular regions. (**F**) G-SMAP analysis. In the computation process in (E), the local K-function and its variants, including G_0_ p_i_(r), were calculated for all the sample and random simulation data points. G-SMAP images were generated by integrating the G_0_ p_i_(r) kernel around each point. Example G-SMAP images for sample and random simulation data in the same ROI are shown in pseudo rainbow colors (upper left panel). A pixel-intensity histogram of the G-SMAP images is shown in the upper right panel. Pixels above a threshold value (e.g., 4 SD of the random distribution) were defined as cluster regions, and closed cluster regions were numbered as individual clusters (pseudo rainbow colors in bottom panels).

**Figure S2.**
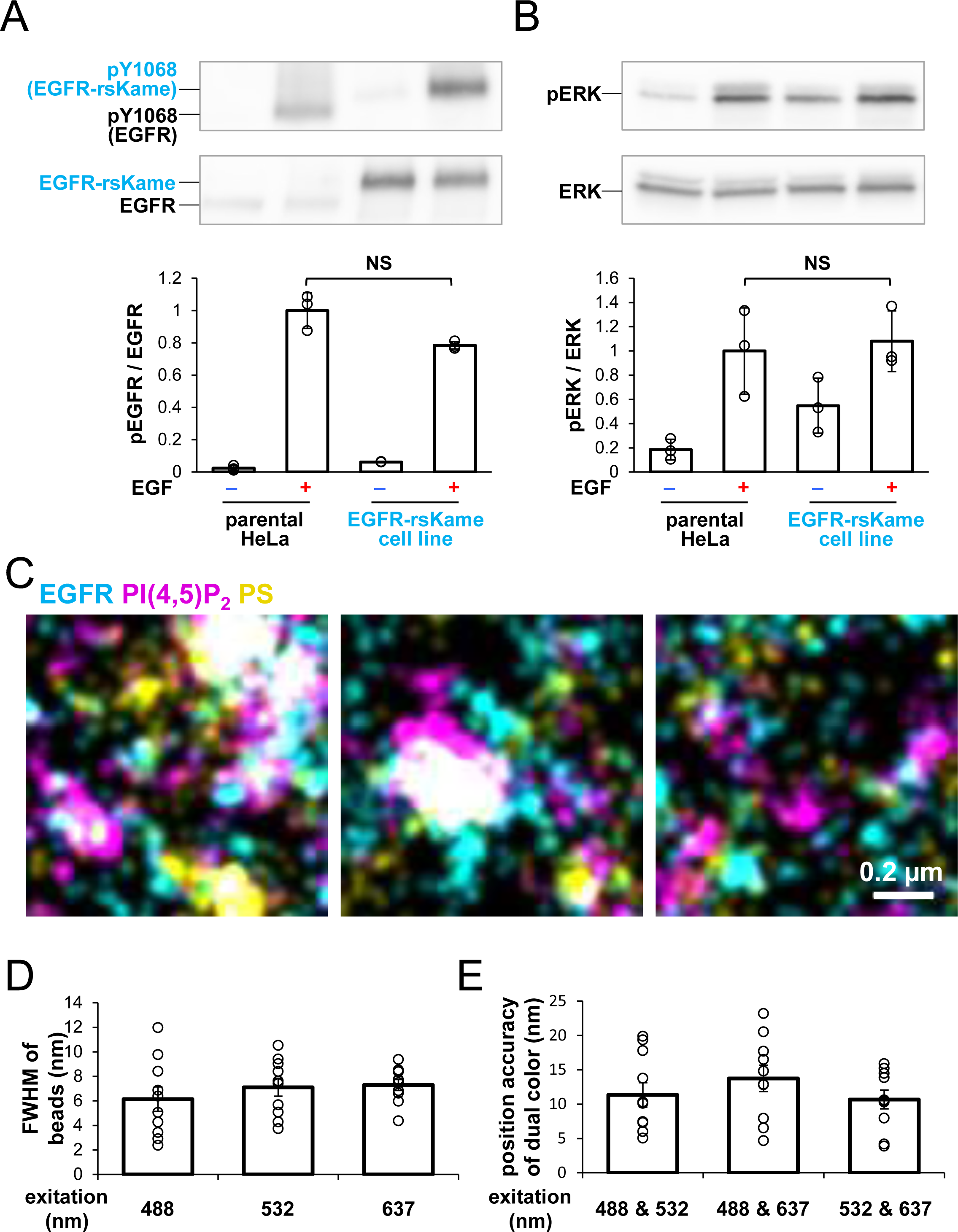
Construction and evaluation of multicolor SMLM. (**A**) Amounts of phosphorylated-Tyr1068 EGFR in the cell line. Top: Western blotting analysis of phosphorylated-Tyr1068 EGFR and total EGFR. Parental HeLa cells and EGFR-KO cells stably expressing EGFR–rsKame fusion protein were incubated in serum-free medium overnight and stimulated with 20 nM EGF for 1 min. Phospho-EGFR and total EGFR were detected with anti-pY1068 EGFR and EGFR, respectively. Bottom: Ratio of phosphorylated-Tyr1068 EGFR/total EGFR. The ratio was normalized to the mean value of parental HeLa cells after EGF stimulation. Data are means ± SD of three experiments. (**B**) Amounts of phosphorylated ERK in the cell line. Top: Western blotting analysis of phosphorylated ERK and total ERK. Cells were prepared as described in (A). Phospho-ERK and total ERK were detected with anti-pERK and ERK, respectively. Bottom: Ratio of phosphorylated-ERK/total ERK. The ratio was normalized to the mean value of parental HeLa cells after EGF stimulation. Data are means ± SD of three experiments. (**C**) Enlarged images of square regions surrounded by thin lines in Fig. 1A. (**D**) Full-width half-maximum (FWHM) of the positional distributions for single immobile fluorescent beads. Data are means ± SEM of 10 beads. (**E**) Position accuracy of the dual-color immobile fluorescent beads. The accuracy of the superimposition of two images was calculated after affine transformation. Data are means ± SEM of 10 beads. NS (not significant) on Welch’s *t* test.

**Figure S3.**
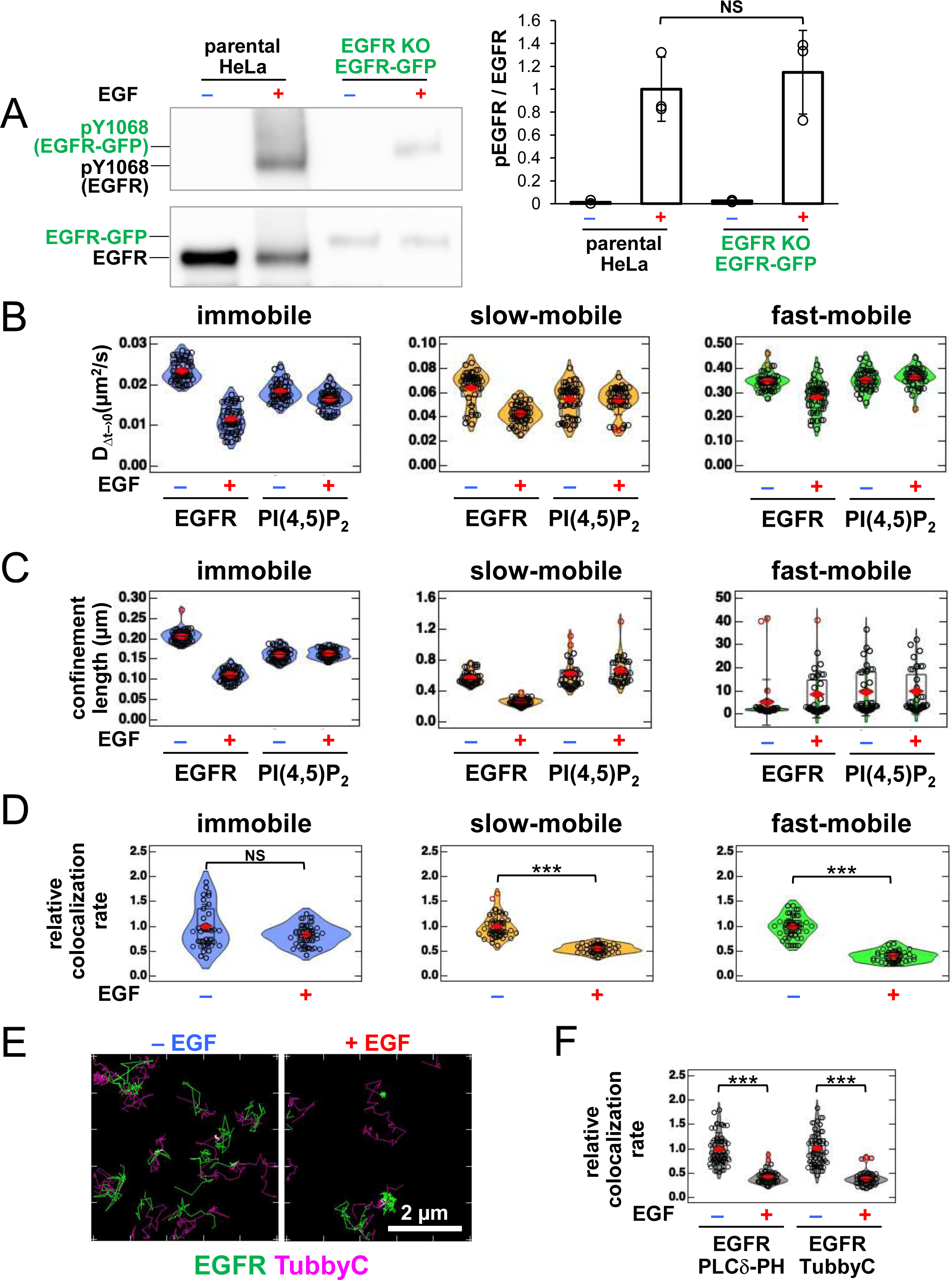
SMT analysis of EGFR–EGFP and JF549–PI(4,5)P_2_. (**A**) Amounts of phosphorylated-Tyr1068 EGFR in the cell for SMT analysis. Left: Western blotting analysis of phosphorylated-Tyr1068 EGFR and total EGFR. Parental HeLa cells and EGFR-KO cells transiently expressing EGFR–GFP at low levels were incubated in serum-free medium overnight and stimulated with 20 nM EGF for 1 min. Phospho-EGFR and total EGFR were detected with anti-pY1068 EGFR and EGFR, respectively. Right: Ratio of phosphorylated-Tyr1068 EGFR/total EGFR. The ratio was normalized to the mean value of parental HeLa cells after EGF stimulation. Data are means ± SD of three experiments. (**B**) Diffusion coefficients of EGFR–EGFP and JF549–PI(4,5)P_2_. Cells were prepared as described in Fig. 3A. (**C**) Confinement lengths of EGFR–EGFP and JF549–PI(4,5)P_2_. (**D**) Relative colocalization rate of EGFR–EGFP and JF549–PI(4,5)P_2_ in the immobile (blue), slow-mobile (yellow), and fast-mobile (green) fractions before and after incubation with 20 nM EGF. To consider the differences in expression levels among cells, the colocalization rate of each fraction was divided by the densities of EGFR–GFP and JF549–PI(4,5)P_2_ of the total fractions and normalized to the mean value obtained before EGF stimulation. Violin plots show the mean value and distribution of 30 cells. (**E**) Trajectories of EGFR–GFP (green), JF549–TubbyC (magenta), and colocalization (white) for 5 s before (left) and after incubation with 20 nM EGF (right) in the same cell. EGFR–GFP and Halo–TubbyC were transiently expressed in EGFR-KO HeLa cells. After overnight incubation in serum-free medium, Halo–TubbyC was stained with JF549–Halo ligand. EGFR–GFP and JF549– TubbyC were observed at a time resolution of 30 ms for 5 s before EGF stimulation. After stimulation with 20 nM EGF, EGFR–GFP and JF549–TubbyC were observed in the same cells between 1 and 5 min after the addition of EGF. (**F**) Relative colocalization rate of EGFR and biosensors of PI(4,5)P_2_ before and after incubation with 20 nM EGF. To consider the differences in expression levels among cells, the colocalization rate was divided by the densities of EGFR– GFP and biosensors of PI(4,5)P_2_, and normalized to the mean value of EGFR–GFP and JF549– PLCδ–PH (JF549–PI(4,5)P_2_) obtained before EGF stimulation. Violin plots show the mean value and distribution of 36 cells. ***p < 0.001, NS (not significant) on Welch’s t test.

**Figure S4.**
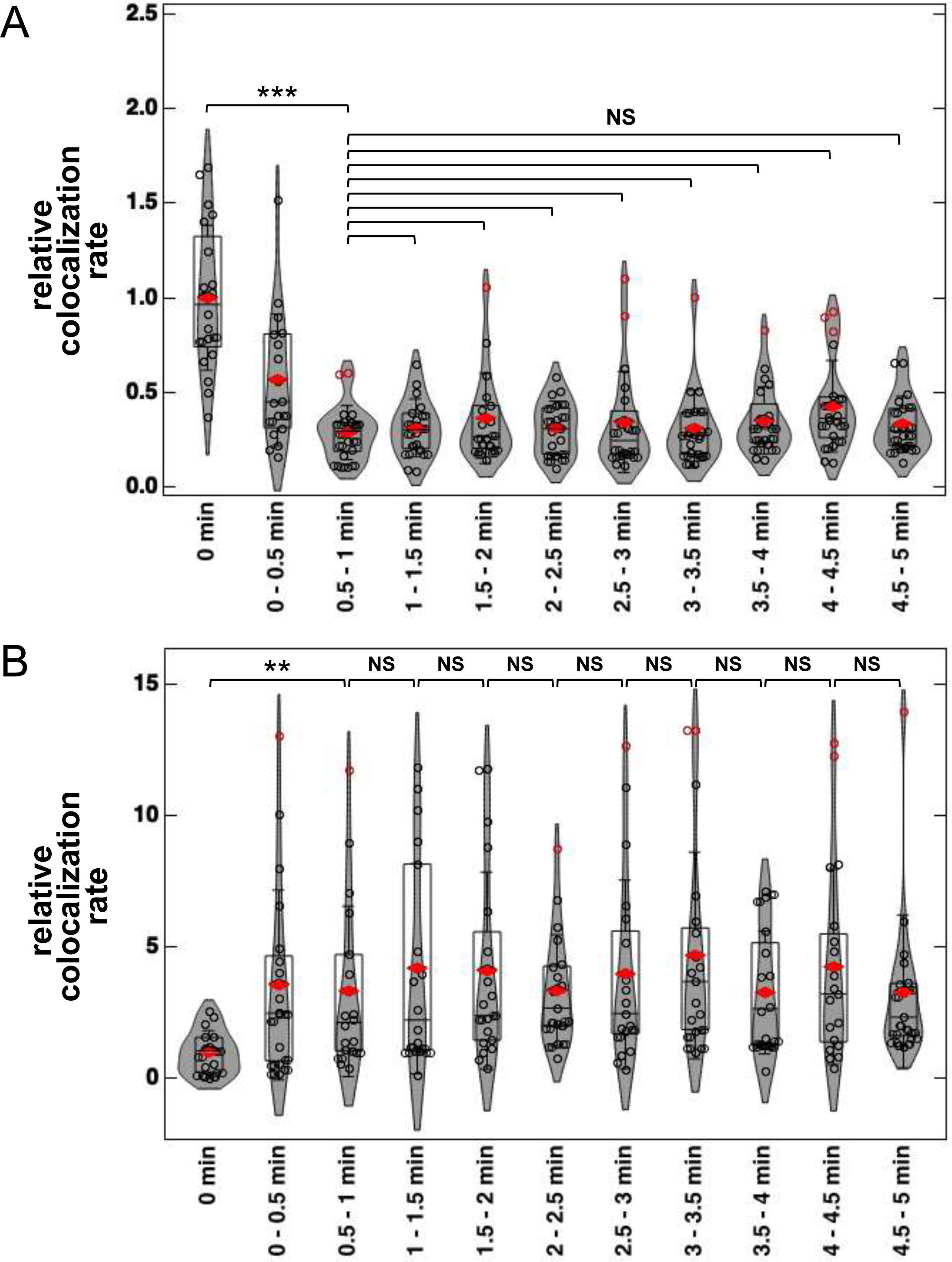
Time-dependent colocalization of EGFR-PI(4,5)P_2_ and EGFR-GRB2. (**A**) Relative colocalization rate of EGFR–GFP and JF549–PI(4,5)P_2_ before and after incubation with 20 nM EGF. EGFR–GFP and Halo–PI(4,5)P_2_ were transiently expressed in EGFR-KO HeLa cells. After the cells were incubated in serum-free medium overnight, Halo–PI(4,5)P_2_ was stained with JF549–Halo ligand. EGFR–GFP and JF549–PI(4,5)P_2_ were observed in different cells from 0 to 5 min after EGF stimulation. To consider the differences in expression levels among cells, the colocalization rate was divided by the densities of EGFR–GFP and JF549– PI(4,5)P_2_ and normalized to the mean value obtained before EGF stimulation. Violin plots show the mean value and distribution of the 18–20 cells. ***p < 0.001, NS (not significant) on Tukey’s multiple comparison test. (**B**) Relative colocalization rate of EGFR–GFP and JF549–GRB2 before and after incubation with 20 nM EGF. EGFR–GFP and Halo–GRB2 were transiently expressed in EGFR-KO HeLa cells. Similar to EGFR–GFP and Halo–PI(4,5)P_2_ in Fig. S4A, EGFR–GFP and JF549–GRB2 were observed in different cells from 0 to 5 min after EGF stimulation. To consider the differences in expression levels among cells, the colocalization rate was divided by the densities of EGFR–GFP and JF549–GRB2 and normalized to the mean value obtained before EGF stimulation. Violin plots show the mean value and distribution of the 17–20 cells. **p < 0.01, NS (not significant) on Welch’s t test.

**Figure S5.**
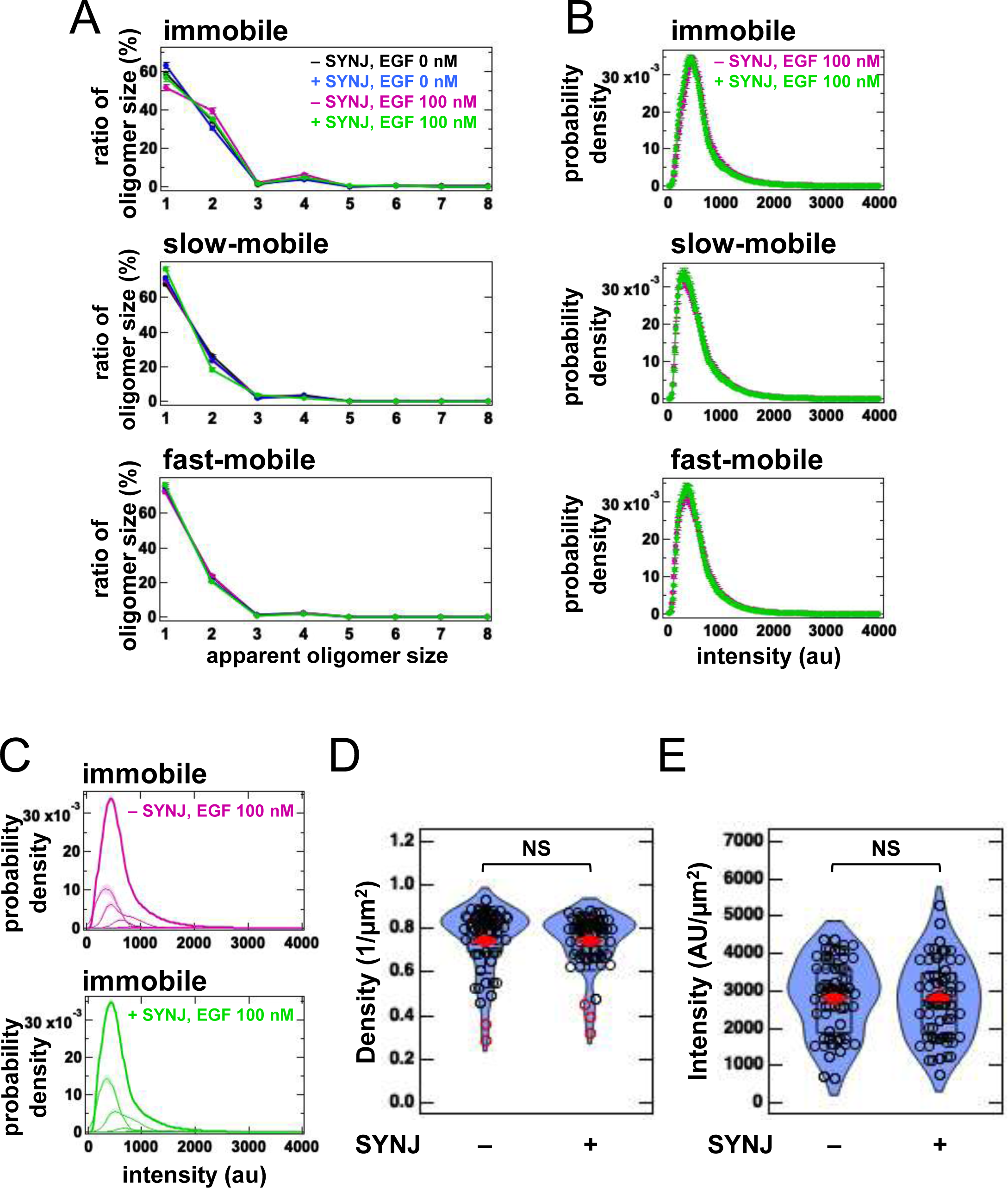
SMT analysis of EGFR under PI(4,5)P_2_-depleted condition. (**A**) SMT analysis of EGFR–SF650 in control and SYNJ-expressing HeLa cells. EGFR–Halo was transiently expressed in EGFR-KO HeLa cells. After cells were incubated in serum-free medium overnight, EGFR–Halo was stained with the SF650–Halo ligand. EGFR–SF650 was observed in control (n = 40) and SYNJ-expressing cells (n = 40) at a time resolution of 30 ms before EGF stimulation. Following stimulation with 100 nM EGF, EGFR–SF650 was observed in the same cells between 1 and 6 min after the addition of EGF. (**B**) Distribution of the total fluorescence intensity of EGFR–SF650 in control (magenta) and SYNJ-expressing cells (green) after stimulation with 100 nM EGF in immobile (top), slow-mobile (middle), and fast-mobile fractions (bottom). EGFR–SF650 in control (n = 40) or SYNJ-expressing cells (n = 40) was observed as described in Fig. 4A. Data are means ± SEM. (**C**) Distribution of the fluorescence intensity of EGFR–SF650 in control (magenta) and SYNJ-expressing cells (green) after stimulation with 100 nM EGF in the immobile fraction. The putative oligomer size was estimated from the total fluorescence intensity (bold line) with Gaussian functions (thin lines). The estimates are shown in Fig. 4A. (**D**) The mean density of EGFR–SF650 in control (n = 40) and SYNJ-expressing cells (n = 40) before EGF stimulation. (**E**) The mean intensity of EGFR–SF650 in control (n = 40) and SYNJ-expressing cells (n = 40) before EGF stimulation. The intensities of EGFR–SF650 were measured at the first frame of the observation and normalized with the area. Violin plots show the mean value and distribution of 40 cells. NS (not significant) on Welch’s *t* test.

**Figure S6.**
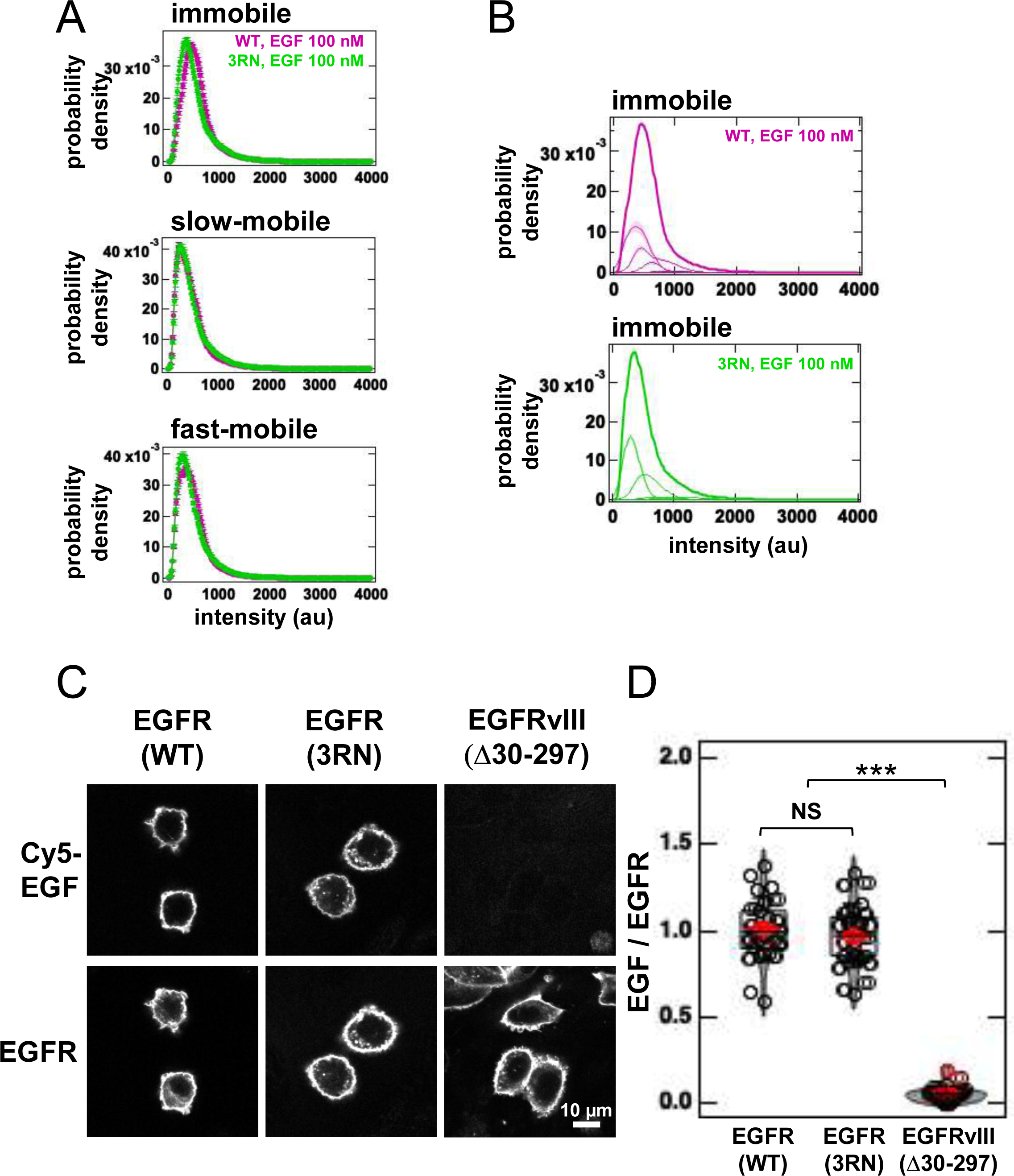
SMT analysis of EGFR under PI(4,5)P_2_-interaction-depleted condition. (**A**) Distribution of the total fluorescence intensity of EGFR(WT)–SF650 (magenta) and EGFR(3RN)–SF650 (green) after stimulation with 100 nM EGF in immobile (top), slow-mobile (middle), and fast-mobile fractions (bottom). EGFR(WT)–SF650 (n = 40) and EGFR(3RN)– SF650 (n = 45) were observed as described in Fig. 4B. Data are means ± SEM. (**B**) Distribution of the fluorescence intensities of EGFR(WT)–SF650 (magenta) and EGFR(3RN)–SF650 (green) after stimulation with 100 nM EGF in the immobile fraction. The putative oligomer size was estimated from the total fluorescence intensity (bold line) with Gaussian functions (thin lines). The estimates are shown in Fig. 4B. (**C**) Images of EGFR (top) and Cy5-labeled EGF (bottom). EGFR(WT), EGFR(3RN), or EGFRvIII lacking amino acid resides involved in the ligand binding in the extracellular region was expressed in EGFR-KO cells and labeled with Cy5-EGF for 10 min. (**D**) The relative intensity of Cy5–EGF bound to EGFR. To consider the difference in the expression level of EGFR, the fluorescent intensities of Cy5-EGF and EGFR–JF549 were measured, and the intensity of Cy5-EGF was normalized with that of EGFR–JF549. The ratio was normalized to the mean value of control cells. Violin plots show the mean value and distribution of 26 imeges. ***p < 0.001, NS (not significant) on Tukey’s multiple comparison test.

**Figure S7.**
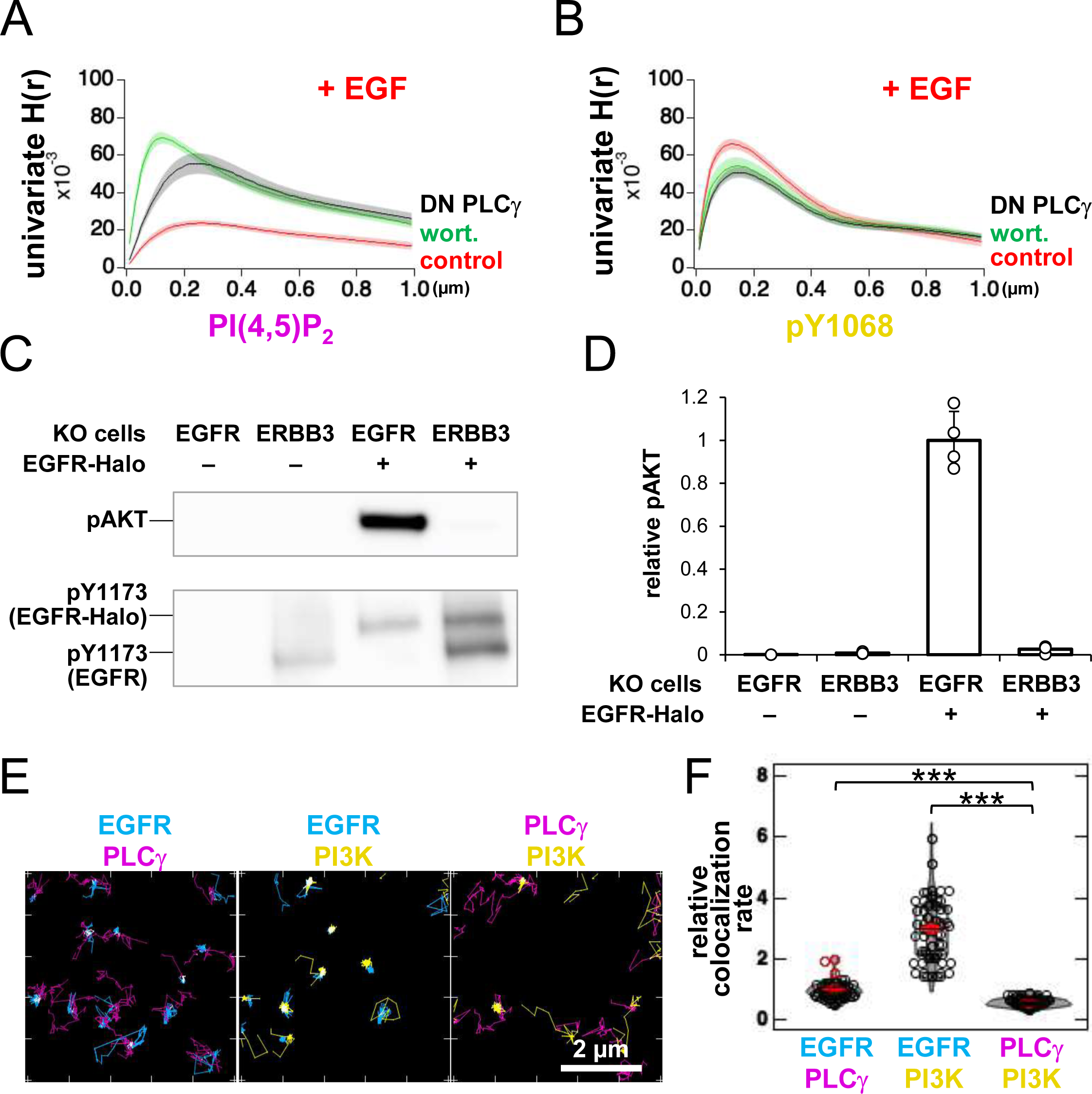
PI3K is not responsible for EGFR dissociation from PI(4,5)P_2_ nanodomains after EGFR is phosphorylated. (**A**) Univariate H(r) value of PAmCherry–PI(4,5)P_2_ in control (red), wortmannin-treated (green), and DN PLCγ-expressing cells (black) after incubation with 20 nM EGF for 1 min. Cells were prepared as described in Fig. 6D. Data are means ± SEM of nine cells. (**B**) Univariate H(r) value of HMSiR–pY1068 in control (red), wortmannin-treated (green), and DN-PLCγ-expressing cells (black) after incubation with 20 nM EGF for 1 min. Cells were prepared as described in Fig. 6D. Data are means ± SEM of nine (control) or 10 (wortmannin-treated and DN PLCγ-expressing) cells. (**C**) Western blotting analysis of phosphorylated AKT (top) and phosphorylated EGFR (bottom). After EGFR-KO and ERBB3-KO cells were transfected with EGFR–Halo, the cells were incubated in serum-free medium overnight and stimulated with 20 nM EGF for 2 min. Phosphorylated AKT and phosphorylated EGFR were detected with anti-phospho-AKT (Ser473) and anti-pY1173 EGFR antibodies, respectively. (**D**) Amount of phosphorylated AKT. Relative intensities were normalized to the mean of lane #3 in (C). Data are means ± SD of four experiments. (**E**) Trajectories of EGFR–GFP (cyan), PLCγ–TMR (magenta), SF650–PI3K (yellow), and colocalization (white) for 6 s after incubation with 20 nM EGF (right). EGFR–GFP, PLCγ (PLCG1)–SNAP, and Halo–PI3K (p85α) were transiently expressed in EGFR-KO HeLa cells. After overnight incubation in serum-free medium, the cells were stained with TMR–SNAP and SF650–Halo ligands. EGFR–GFP, PLCγ–TMR, SF650–PI3K were observed at a time resolution of 30 ms for 5 s, between 1 and 5 min after the addition of EGF. (**F**) Relative colocalization of EGFR, PLCγ, and PI3K after incubation with 20 nM EGF. To consider the differences in expression levels among cells, the colocalization rate was divided by the density and normalized to the mean value of EGFR–GFP and PLCγ–TMR. Violin plots show the mean value and distribution of 35 cells. ***p < 0.001 based on Tukey’s multiple comparison test.

## Notes

### Competing Interest Statement

The authors have declared no competing interest.

### Summary of Updates

Figures 2B, 2C, and 2D revised. Figure 3E revised. Figure 6E revised. Figures S3A, S3D, S3E, and S3F revised. Figures S4A and S4B revised. Figures S7E and S7F revised.

